# Germline regulation of tumor evolutionary dynamics shapes multiple myeloma progression

**DOI:** 10.64898/2026.06.05.730550

**Authors:** Hai Chen, Jingmin Shu, Rekha Mudappathi, Panwen Wang, Leif Bergsagel, Ping Yang, Zhifu Sun, Changxin Shi, Li Liu

## Abstract

Germline variation shapes cancer risk, yet its influence on the evolutionary dynamics of established tumors remains poorly understood. In multiple myeloma, subclonal diversification drives disease progression and treatment failure, but the heritable factors that modulate this process are unknown.

Here, we show that germline variation is associated with tumor evolutionary features, implicating inherited regulation in subclonal expansion. Integrating germline variation with tumor evolutionary parameters identifies variants associated with evolutionary features, with signals enriched in regulatory regions, consistent with a transcriptional basis. We further identify *TBKBP1* as a key locus linking germline variation to tumor evolution and clinical outcome.

Germline variation at this locus is associated with *TBKBP1* expression and subclonal expansion, and *TBKBP1* expression correlates with adverse prognosis, consistent across independent cohorts. Functional analyses demonstrate that *TBKBP1* promotes proliferation and activates MYC, mTORC1 and non-canonical NF-κB signaling pathway.

Together, these findings establish germline regulatory variation as a determinant of tumor evolutionary dynamics and identify *TBKBP1* as a mediator linking inherited variation to subclonal expansion and disease progression in multiple myeloma.

## INTRODUCTION

Intratumoral heterogeneity, characterized by the coexistence of genetically and phenotypically distinct subclones, poses a major barrier to accurate diagnosis and durable responses to precision therapies^1,2^. Underlying these challenges is the highly dynamic tumor evolutionary process, wherein somatic alterations accumulate and constitute the genetic substrate, upon which natural selection and genetic drift act under microenvironmental constraints, resulting in continual diversification and competitive clonal expansion^3^. Because somatic mutations occur within the context of an inherited germline genome, inter-individual differences in germline variation may modulate the selective landscape and, consequently, the trajectory of tumor evolution^4,5^.

Emerging evidence of germline-somatic interactions (GSI) in cancer has shown that somatic mutational features^6–9^, including mutation burden, mutational signatures, and the prevalence of specific alterations, are associated with inherited germline variation, both at the level of individual locus and in aggregate (e.g., polygenic risk scores and genetic ancestry). However, these investigations have predominantly focused on static genomic endpoints, such as mutation frequency at the time of biopsy, and do not capture the inherently dynamic nature of tumor development. It remains unresolved whether the germline background influences merely the identity of somatic alterations or also shapes the evolutionary trajectory of a tumor.

In this study, we investigated GSI in an evolutionary framework. Our previous work has shown that the temporal and selective forces governing tumor progression can be quantitatively described by a set of evolutionary parameters inferred from single-timepoint sequencing data^10^,including the mutation rate, subclone fitness, subclone emergence time, subclone expansion score, and fitness diversity index. By applying these metrics to a large cohort of multiple myeloma (MM) patients^11^, we provide the first evidence that inherited germline variants predispose tumors to specific evolutionary trajectories. We further show that tumor evolution trajectories may mediate the effects of germline variation on clinical outcome. This integrative analysis identified several novel germline loci associated with MM progression. Via functional assays, we validated a cis-acting locus that upregulates *TBKBP1* expression and subsequently promotes tumor growth, revealing a new mechanism through which inherited variation influences cancer progression.

## RESULTS

### Patient and Sample Characteristics

We analyzed a cohort of 869 primary MM tumors from the CoMMpass study^11^, leveraging both high-depth whole-exome sequencing (WES) (mean coverage 125×) and low-pass whole-genome sequencing (WGS) (mean 6×) data. Using the GDC Data Portal, we retrieved clinical information and annotated somatic mutations detected from the WES of these tumors. The WGS data from matched normal samples were obtained via dbGaP (phs000748.v7.p4.c2). To mitigate the risk of false positives inherent in low-pass sequencing, we applied stringent quality filters via the Mayo Clinic pipeline (**Supplement Methods**). To ensure high-resolution subclonal reconstruction, we used the high-depth WES data and applied TEATIME to infer a suite of quantitative parameters from the somatic mutation landscape. Excluding cases where the TEATIME optimization failed to converge, primarily due to less mutation clustering and low mutation burden (Supplementary Fig. 1A & B), a final cohort of 418 tumors was retained for downstream GSI and clinical association analyses (**Methods**).

Table 1 summarizes the characteristics of the 418 patients. The cohort is representative of the general demographical distribution of MM in the United States, with a mean age at diagnosis of 63.6 years (range, 27 – 89) and a slight male predominance (57% versus 43%). Clinical staging at diagnosis was nearly evenly distributed across the three major stages (31.1%–34.7%), capturing a broad spectrum of disease severity and progression. The somatic mutational landscape, derived from high-depth WES, exhibited a mean Tumor Mutational Burden (TMB) of 1.29 (log(mutations/Mb)) and standard deviation (SD) of 0.55. An average of 43301 single-nucleotide variants (SNVs) were identified per germline genome, and the population allele frequency distribution was skewed toward low-frequency variants, with a median of 0.188 (Supplementary Fig. 2). This relatively low variant yield reflects the conservative filtering criteria applied and the low-pass WGS data.

We applied TEATIME to estimate evolutionary parameters of individual tumors. TEATIME models a tumor as a mixture of two competing cell populations: an ancestral clone with baseline fitness and a derived subclone with elevated fitness, and estimates mutation rates (µ), subclone emergence time (*t*_*f*_), subclone fitness (s), subclone expansion score (τ), and tumor fitness diversity (θ). For the 418 primary tumors analyzed, these parameters exhibited substantial inter-tumor variability (**Fig. 1**). The median mutation rate is 4.5 mutation per cell division (IQR, 3.57–5.78). The fitness diversity index (θ) had a median of 0.95 (IQR, 0.9-0.98). The subclone fitness (s) showed a median of 0.68 (IQR, 0.31–0.90), subclone emergence time (*t*_*f*_) had a median of 4.0 (IQR, 2.3–5.0), and the subclone expansion score (τ) had a median of 2.8 (IQR, 2.25–4.72). UMAP embedding of all clonal evolution parameters showed no separation by gender or ISS stage, suggesting these phenotypes are not confounded by demographic or clinical staging variables. We also examined whether tumor evolutionary parameters were associated with patient demographic and clinical characteristics independently, and observed no significant differences between sex, age, or clinical stages groups (**Supplementary Figure 3**), indicating broadly comparable evolutionary dynamics across baseline clinical strata.

**Table 1.** Summary of patient characteristics.

| Characteristics | Summary Statistics |
| --- | --- |
| Age at diagnosis (years) | Mean: 63.6, SD: 10.1 |
| Sex | Male: 57%, Female 43% |
| Clinical stage | Stage I: 34.2%, II: 34.7%, III: 31. |
| Germline variants per sample # | Mean: 43304.2, SD: 1616.1 |
| Tumor mutational burden (TMB, ln(mutations/Mb)) | Mean: 1.29, SD: 0.55 |

**Figure 1.**
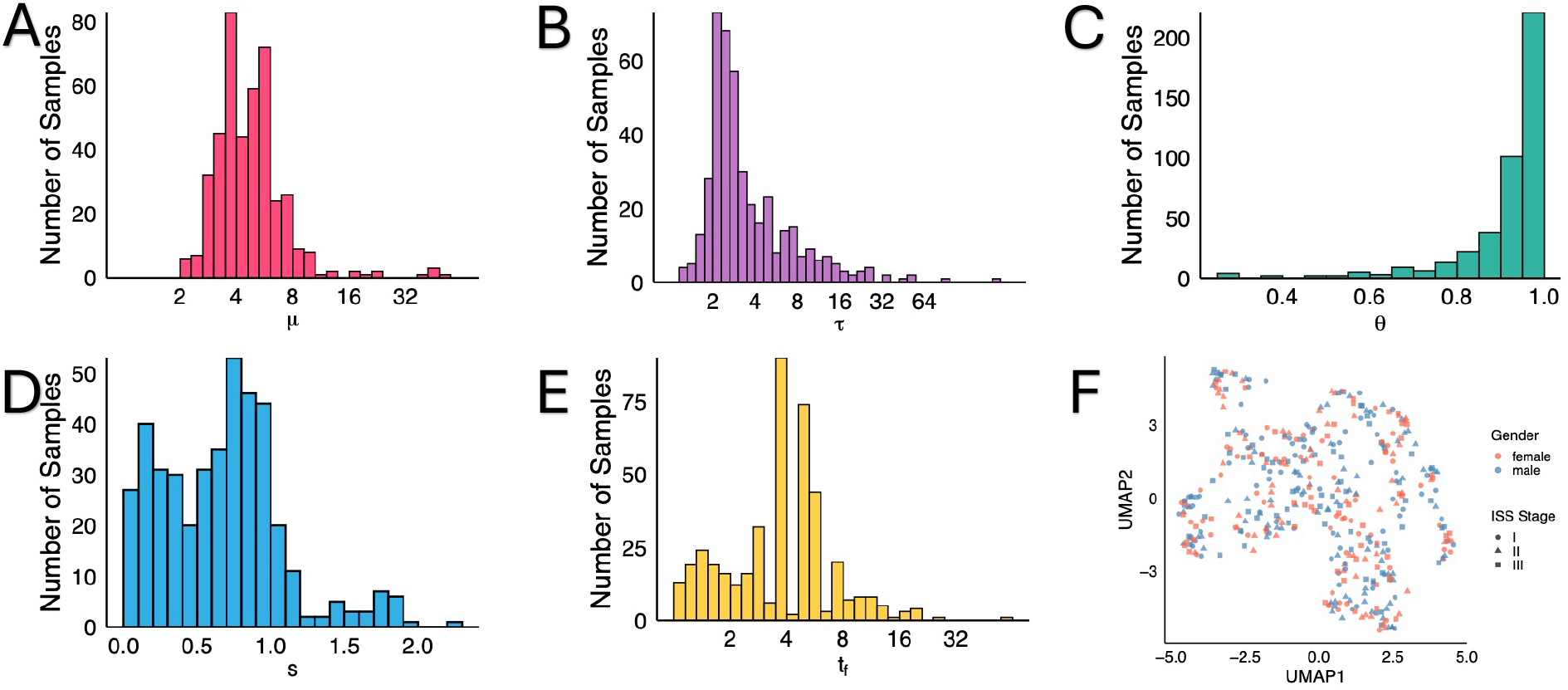
Characteristics of evolutionary parameters. **(A-E).** The mutation rate (μ), subclone emergence time (*t*_*f*_) and subclone expansion score (τ) are log2 transformed to facilitate visualization. (**F**). UMAP embedding of all clonal evolution parameters.

### Evolutionary Dynamics are Associated with Patient Overall Survival

We next assessed the impact of tumor evolutionary dynamics and patient overall survival (OS). To account for the potential interdependence of these metrics, we fitted a multivariable Cox proportional hazards model that included all estimated evolutionary parameters simultaneously, adjusting for age, sex, TBM and karyotype (**Table 2**). In the full cohort, an elevated *τ* score, reflecting a more prolonged period of subclone expansion, was significantly associated with shorter OS (Hazard ratio HR = 1.02, *P* = 0.036).

**Table 2.** Survival analysis for various tumor stages.

|  | Stage I |  | Stage II |  | Stage III |  | All Stage |  |
| --- | --- | --- | --- | --- | --- | --- | --- | --- |
|  | HR | *P* | HR | *P* | HR | *P* | HR | *P* |
| $\mu$ | 0.79 | 0.300 | 0.85 | 0.130 | 1.13 | 0.030 * | 0.96 | 0.300 |
| $\tau$ | 0.98 | 0.900 | 1.02 | 0.120 | 1.07 | 0.046 * | 1.02 | 0.036 * |
| $\theta$ | 1.41 | 0.500 | 1.08 | 0.700 | 0.96 | 0.800 | 1.02 | 0.800 |
| $t_f$ | 0.77 | 0.300 | 1.01 | 0.900 | 0.98 | 0.700 | 1.00 | 0.900 |
| $s$ | 0.13 | 0.200 | 1.15 | 0.800 | 0.97 | 0.900 | 0.84 | 0.600 |
| Age | 1.01 | 0.800 | 1.04 | 0.064 † | 1.03 | 0.075 † | 1.04 | 0.001 *** |
| Sex (Male) | 7.99 | 0.059 † | 2.18 | 0.052 † | 0.87 | 0.600 † | 1.66 | 0.026 * |
| TMB | 7.56 | 0.043 * | 2.79 | 0.100 † | 0.67 | 0.500 | 1.72 | 0.075 † |
| Karyotype | 4.8 | 0.071 † | 0.51 | 0.086 † | 0.87 | 0.700 | 0.81 | 0.300 |

HR: Hazard ratio. †*P* ≤ 0.1; \* *P* ≤ 0.05; \*\* *P* ≤ 0.01; \*\*\* *P* ≤ 0.001

Interestingly, stratification by clinical stage showed that none of the evolutionary parameters were significantly associated with survival in Stage I or II tumors. In contrast, among stage III tumors, the subclone expansion score remained a robust predictor of poorer survival (HR = 1.07, *P* = 0.046). Additionally, a higher mutation rate (*μ*) emerged as a significant independent risk factor for worse overall survival in this subgroup (HR = 1.13, *P* = 0.03). These results suggest that the clinical impact of evolutionary trajectories is most pronounced in advanced-stage disease, where a high intrinsic mutation rate and the sustained expansion of competitively fit subclones converge to drive aggressive progression.

### Germline variation modulates high-risk tumor evolutionary dynamics

Because elevated mutation rate and prolonged subclone expansion were significant predictors of poor patient prognosis in Stage III disease, we investigated whether inherited germline variation contributes to these unfavorable evolutionary dynamics. Specifically, we performed a genome-wide association study (GWAS) for each evolutionary parameter. Given a germline variant, we fitted a linear regression model with the evolutionary parameter as the continuous response and the genotype as the predictor, adjusting for age, sex, TBM, karyotype, and top 10 principal components derived from germline variants to account for population structure (**Methods and Materials**). We tested associations under three genetic models, additive, dominant, and recessive, to maximize power for detecting effects across different inheritance patterns. In the recessive model, the homozygous alternative genotype was coded as 1 and all other genotypes were coded as 0; in the dominant model, carriers of at least one alternative allele were coded as 1; and in the additive model, genotype dosage (0, 1, 2) was used directly.

As expected, the overwhelming majority of germline variants showed no significant associations with tumor evolutionary dynamic parameters (*μ* or *τ*, **Fig. 2A-B**), with only a small number of loci exhibiting extremely low *P* values. Quantile–quantile analysis confirmed that the observed *P* values adhered to the expected null distribution, with deviation only in the upper tail, indicating the presence of true association signals rather than systematic inflation (**Fig. 2C, supplement Fig.4**). To account for multiple testing and linkage disequilibrium (LD) structure, we adjusted nominal *P* values to derive LD-level false discovery rates (FDR, **Methods**). In total, 813 independent LD blocks were significantly associated with tumor evolutionary parameters (FDR <0.05, **Supplementary Table 1)**, hereafter referred to as GSI loci. 17 of these GSI loci exhibited pleiotropic effects, showing significant associations with both *μ* and *τ* (**Fig. 2D)**.

**Figure 2.**
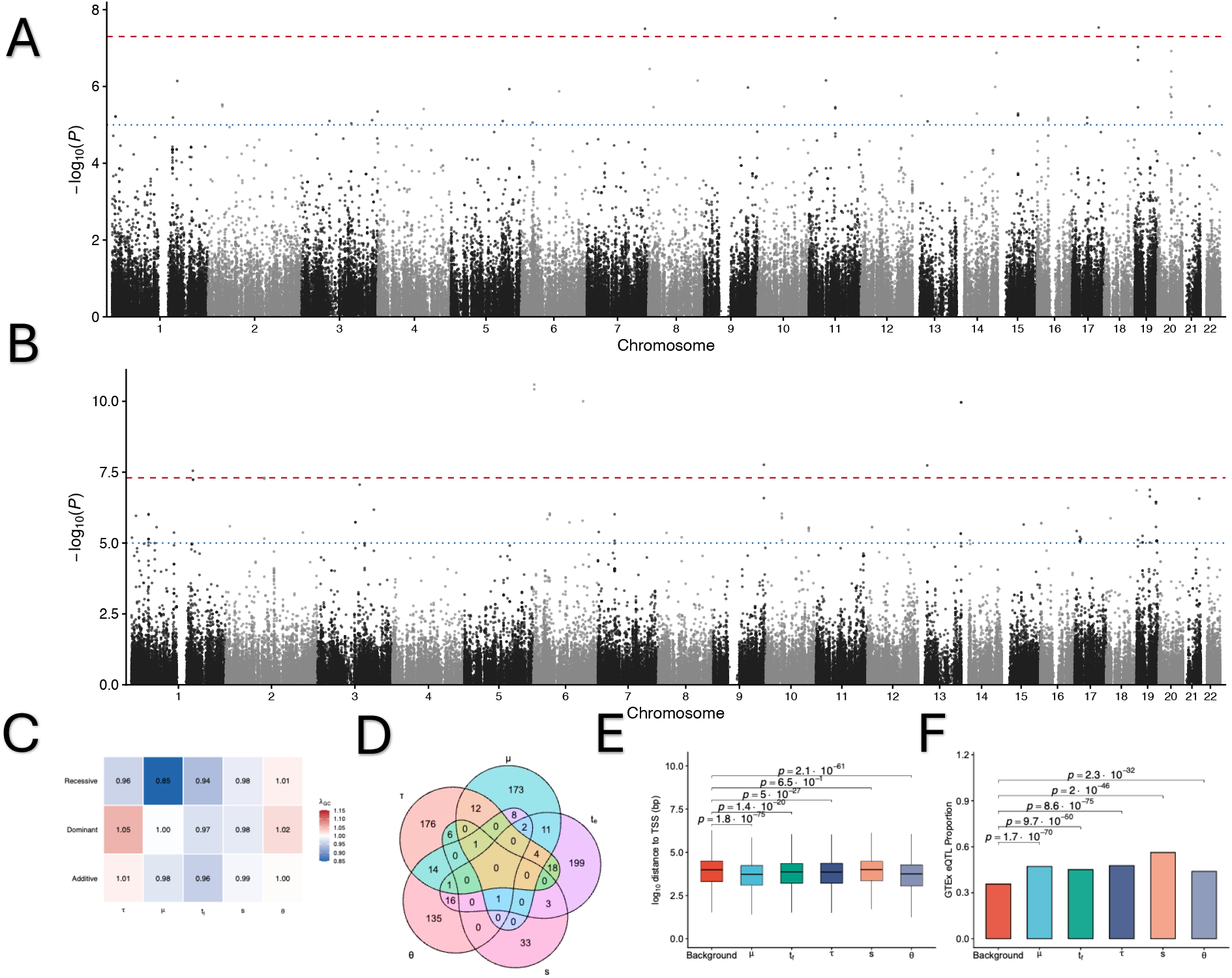
Genome wide association test results. (**A-B**) Manhattan plot of association test results for mutation rate (A) and subclone expansion score (B). (**C**) Genomic inflation factors across five evolutionary parameters and three genetic models. (**D**) Venn diagram showing the overlap of loci with significant associations with mutation rate and subclone expansion score. (**E**) Distance between germline variants with significant associations to transcription start sites. (**F**) Proportion of loci overlapping with eQTLs.

Among the 17 pleiotropic loci, several harbored genes involved in lymphocyte survival and innate immune signaling, including *TNFRSF13B, GIMAP5, CD101*, and *SIGLEC5/14*. These dual associations suggest germline variants at these loci may modulate fundamental mechanisms coordinating both mutational influx and subsequent clonal outgrowth.

To assess the functional potential of the identified GSI loci, we characterized their genomic distribution relative to transcription start sites (TSS)^12^ and overlap with expression quantitative trait loci^13^ (eQTLs, **Supplement Methods**). Compared to non-GSI loci, GSI loci were significantly closer to TSS (t-test *P* = 8.1×10^−85^, **Fig. 2D, Supplement Fig 5A)** and exhibited a markedly higher proportion of eQTLs reported in the normal whole blood samples in the GTEx study (χ^2^ test *P* = 2.8×10^−207^, respectively, **Fig. 2E, Supplement Fig 5B**). This enrichment in regulatory regions suggests that the germline influence on tumor evolutionary trajectory is primarily mediated via transcriptional modulation.

### GSI is associated with patient overall survival

We next evaluated the clinical relevance of the identified GSI loci. Cox proportional hazard models revealed that 1026 SNVs across 228 loci were significantly associated with patient overall survival (OS). To assess whether the inherited risk for poor prognosis is realized through the pre-programming of somatic evolutionary trajectories, we performed a causal mediation analysis^14^ and identified 90 SNVs spanning 32 loci with significant indirect effects (FDR <0.05), indicating that their influence on overall survival was mediated through the evolutionary parameters. Of these, all SNVs were mediated through subclone expansion.

Given the regulatory enrichment of associated loci, we next assessed whether variants within these loci modulate the expression of their putative target genes in multiple myeloma (**Supplement Methods**) This analysis identified 8 regulatory GSI loci (7q22.1, 3p22.3, 21q22.3, 1p13.2, 1p34.1, 7p13, 19p13.12, and 17q21.32), encompassing 15 SNVs, for which the genotypes were associated with the expression levels of 9 genes (*PILRB, STAG3L5P, STAC, SIK1, ST7L, PRDX1, LINC00957, CYP4F2*, and *TBKBP1*). To prioritize genes with potential clinical relevance, we further required evidence of prognostic association^15,16^(**Supplement Methods)** and directional consistency across genotype, gene expression, evolutionary parameters and survival outcome, Under these criteria, only three genes, *STAC, LINC00957* and *TBKBP1*, remained supported. All exhibited concordant patterns, in which the allele associated with increased expression of the target gene was also associated with more aggressive tumor evolutionary trajectories and poorer prognosis, suggesting that germline risk may be mediated through transcriptional regulation that subsequently influences tumor evolutionary dynamics and clinical outcomes (**Fig. 3A**). We prioritized *TBKBP1* for its established role in NF-κB signaling^17^, providing a biologically plausible link between germline variants and tumor evolution. The locus is 17q21.32 harboring the *TBKBP1* genes. We analyzed eQTLs within ±500 kb of each gene and examined their effect sizes adjusting for covariate (Fig. 3B, supplementary Fig.6). At the *TBKBP1* locus, eQTL signals coalesce within an upstream LD block that includes variants retained after mediation analysis, whereas downstream signals are more dispersed across smaller, localized LD blocks, indicating a more heterogeneous regulatory architecture.

**Figure 3.**
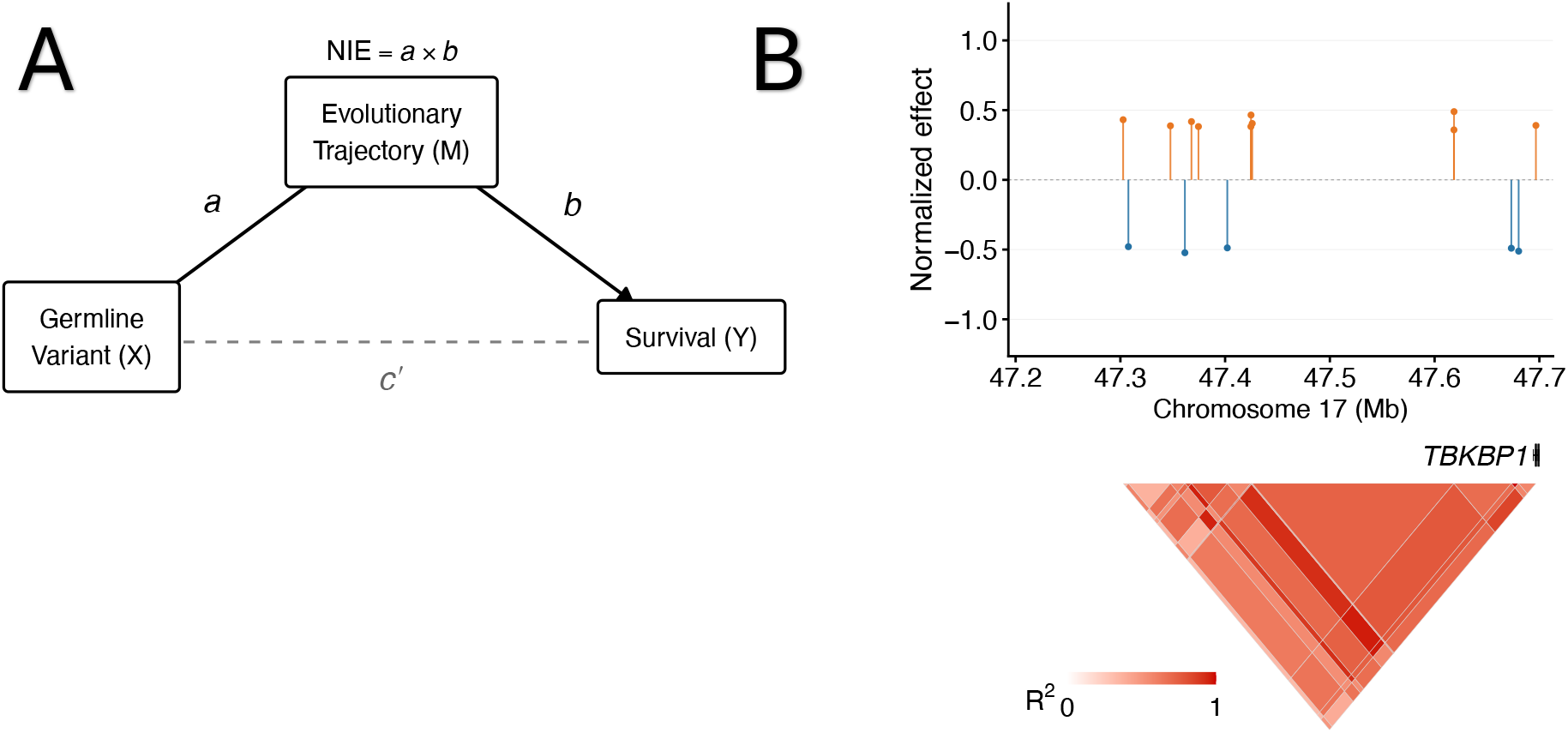
Regulatory enrichment and overlap of germline loci associated with somatic evolutionary parameters. (**A**) Schematic representation of the mechanism, in which genomic variants dysregulate gene expression, which modulate the tumor evolutionary trajectory, and eventually leads to poor clinical outcomes. **(B)** Cohort eQTLs. The top panel shows normalized effect sizes of eQTLs. Gene models are shown in the middle panels. The bottom panel shows local LD structure (R^2^).

### *TBKBP1* expression links germline variation to subclonal dynamics

Among the lead GSI loci, we identified a high-confidence association at the rs8078309 locus. Variants at this locus were significantly associated with increased *TBKBP1* expression and prolonged subclonal outgrowth (**Fig. 4 A-B**, both *P* <0.05). Consistent with our earlier observation that prolonged subclonal expansion is associated with adverse clinical outcomes (**Table 2, Fig. 4C**, *P* =0.0072), we found that higher *TBKBP1* expression was a significant risk factor of poor patient OS (**Fig. 4D**, *P* =0.036). To assess the robustness of this association, we performed external validation in an independent cohort of 554 MM tumors (GSE24080)^18^, which replicated the association between elevated TBKBP1 expression and diminished OS (**Supplementary Fig. 7A**). Together, these findings identify *TBKBP1* as a putative germline-encoded driver of aggressive subclonal dynamics in multiple myeloma.

**Figure 4.**
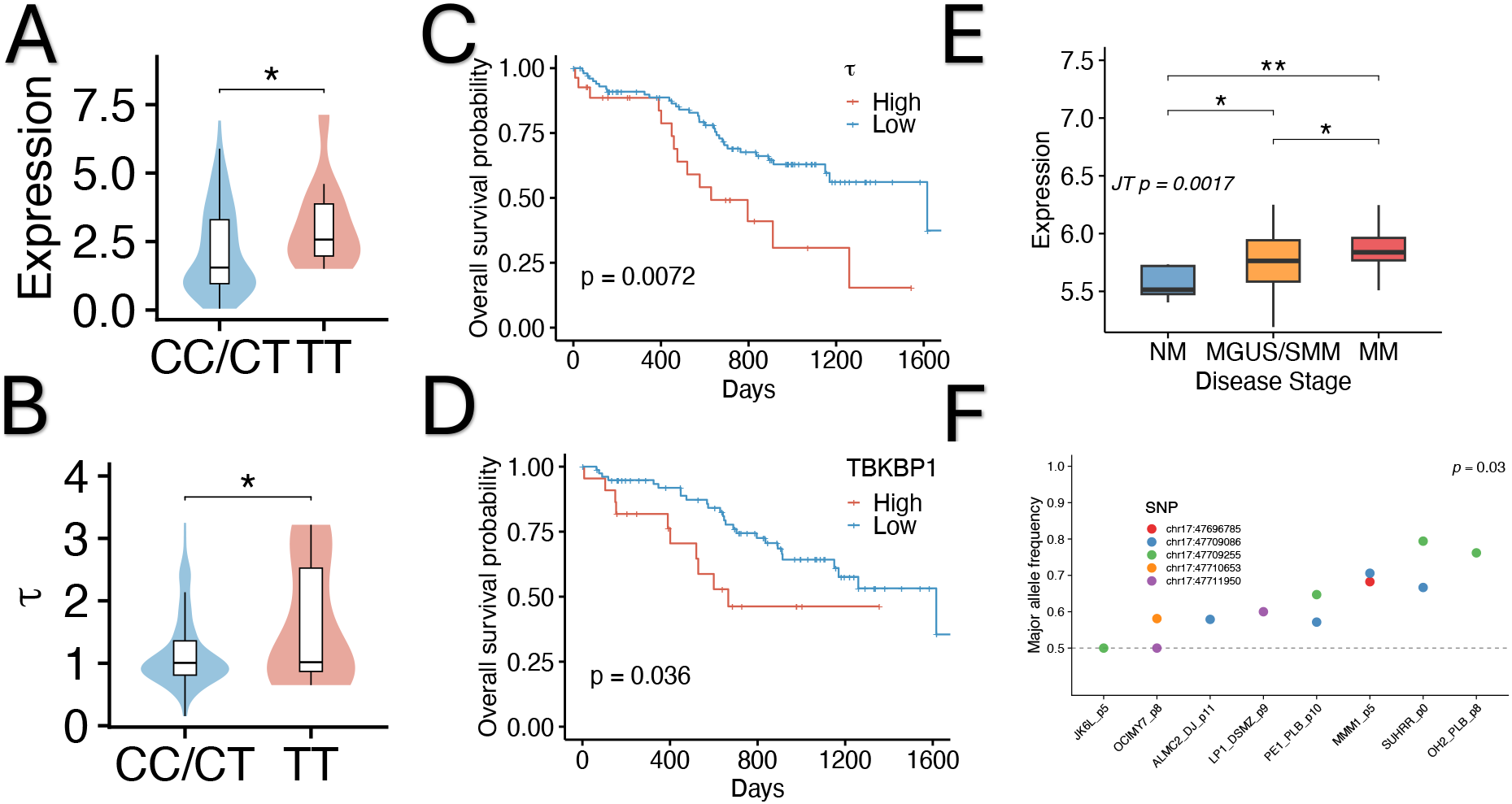
Germline variants at the *TBKBP1* locus associate with expression and subclonal dynamics. (**A**) *TBKBP1* expression stratified by genotype at the lead mediation variant, asterisk denotes statistical significance. (**B**) *τ* by genotype group. Values are log transformed for visualization. (**C-D**) Kaplan–Meier overall survival curves for MMRF stage III patients stratified by *τ* score (C) and by *TBKBP1* expression (D). (**E**) *TBKBP1* expression across disease stages in GSE47552 cohorts. Groups: normal marrow (NM), MGUS/SMM (precursor), and multiple myeloma (MM). (**F**) Major allele frequency per cell line at heterozygous TBKBP1 SNP sites from allele-specific expression analysis (N = 8 cell lines, 5 SNP positions). gene-level P from ASEP nonparametric mixture model (10,000 permutations).

We next examined the role of *TBKBP1* expression in disease progression using two independent cohorts encompassing a total of 20 normal bone marrow, 97 precancerous stages (MGUS/SMM), and 144 MM tumors (GSE47552^19^ and GSE6477^20^). In both cohorts, *TBKBP1* expression increased progressively from normal bone marrow through precursor states to frank malignancy (**Fig. 4E, Supplementary Fig. 7B**, P<0.05). This pattern was supported by a highly significant ordered trend (Jonckheere trend test *P* = 0.0017), with stepwise increases at both the normal-to-precursor and precursor-to-malignant transitions. These observations suggest that *TBKBP1*-associated regulatory activity is engaged early in the myeloma-genesis and becomes progressively more pronounced as the disease advances.

To confirm that the identified germline variants exert direct regulatory effects on *TBKBP1*, we conducted an allele-specific expression (ASE) analysis across eight MM cell lines^21^(**Methods)**. We observed significant allelic imbalance with preferential expression of the *TBKBP1* transcript linked to the risk-associated alleles (**Fig. 4F**, *P* = 0.03). These findings support a cis-regulatory mechanism by which variants at this locus (and linked SNPs in high LD) modulate *TBKBP1* expression, providing a mechanistic link between inherited variation, gene regulation and tumor evolutionary dynamics.

### *TBKBP1* promotes myeloma cell proliferation through MYC and NF-κB signalling

We performed *in vitro* assays to validate the functional consequences of TBKBP1 upregulation. Specifically, we generated stable *TBKBP1*-overexpressing models by transducing KMS11 and MM1.S cell lines with a pCDH-CMV-MCS-EF1-Puro lentiviral vector, followed by puromycin selection to ensure high-efficiency integration. Following transduction, we quantified cell viability over a 20-day incubation period using the MTT assay to measure metabolic activity as a proxy for proliferation (**Methods**). In both models, *TBKBP1* overexpression significantly enhanced cellular fitness, yielding a three-fold higher viability signal compared to empty vector controls by day (**Fig. 5A**). These consistent effects across independent cell lines indicate that TBKBP1 enhances myeloma cell proliferation and fitness in vitro.

**Fig. 5.**
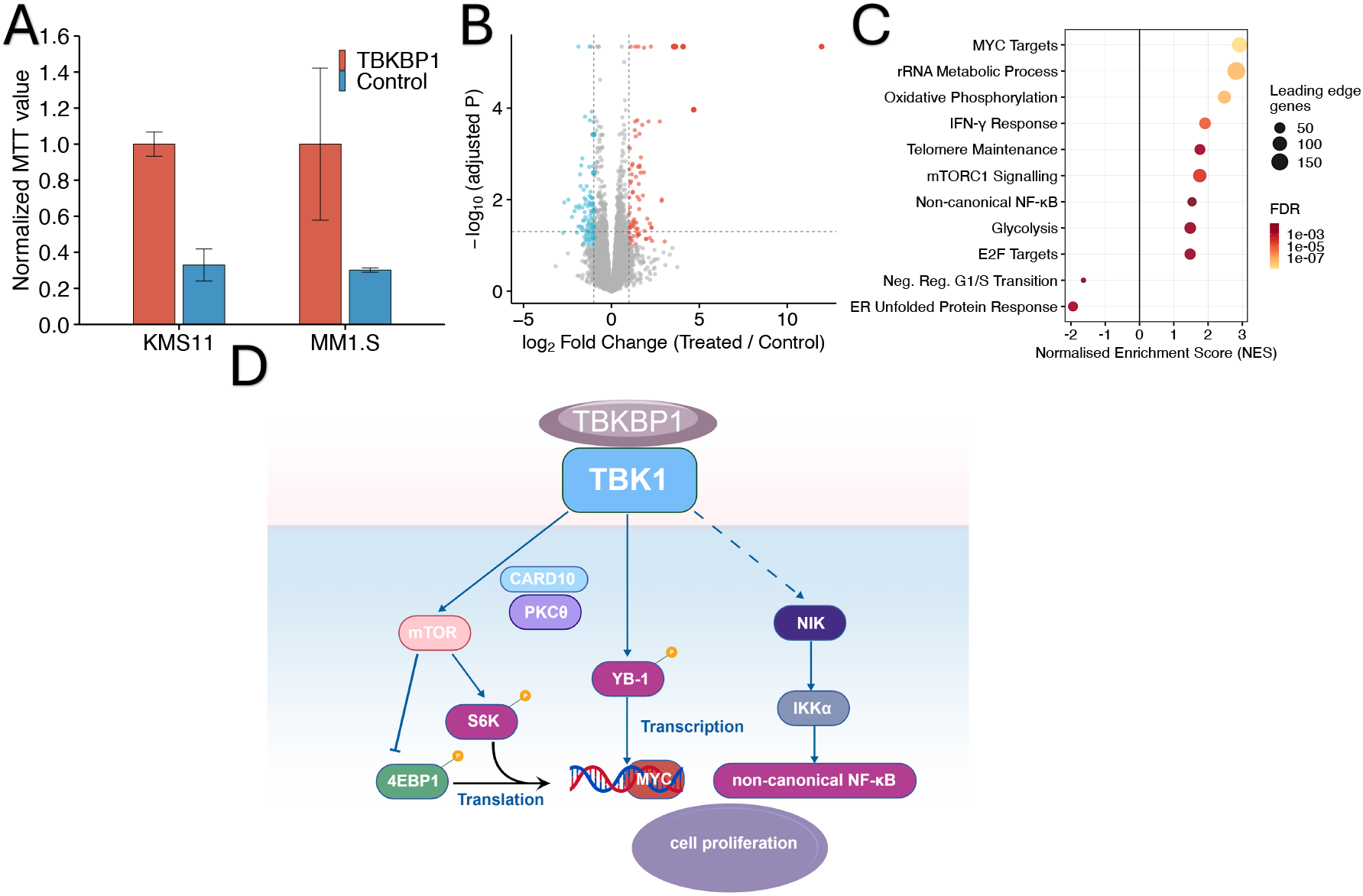
*TBKBP1* overexpression promotes proliferation. (**A**) Normalized MTT cell viability in KMS11 and MM1.S myeloma cell lines expressing *TBKBP1* or empty vector control. Values are normalized to the TBKBP1-overexpressing condition within each cell line. (**B**) Volcano plot showing differential gene expression between *TBKBP1* overexpression and control. Upregulated genes are shown in red and downregulated genes in blue. (**C**) Gene set enrichment analysis (GSEA) plots; NES denotes normalized enrichment score. (**D**) Schematic model illustrating the proposed regulatory network linking *TBKBP1, TBK1* and downstream signaling pathways.

To elucidate the transcriptional mechanisms underlying this enhanced viability phenotype, we performed RNA-seq analysis on *TBKBP1*-overexpressing cells and empty vector controls. A total of 134 gene were significantly differentially expressed (**Fig. 5B**, **Supplementary table 2**). Gene set enrichment analysis (**Fig. 5C**)^22–24^ revealed strong activation of *MYC* targets and *mTORC1* signaling, consistent with accelerated cell-cycle progression^25,26^. Conversely, pathways that restrain proliferation, including the unfolded protein response and negative regulators of the G1/S transition, were significantly suppressed^27^. Metabolic processes, including oxidative phosphorylation and glycolysis, were also upregulated, supporting increased cellular activity^28,29^. Non-canonical NF-κB signaling was also enriched, consistent with its known role in promoting myeloma cell survival and proliferation^30^. These findings indicate that *TBKBP1* promotes myeloma cell proliferation by activating cell-cycle–associated transcriptional processes while relieving constraints on cell-cycle progression.

### A mechanistic model for germline-driven evolution

Integrating these transcriptional findings with the known biochemical functions of *TBKBP1*, we propose a model linking *TBKBP1* upregulation to the proliferative phenotype observed *in vitro* and the elevated *τ* in vivo (**Fig. 5D**)^31^. As an adaptor of *TBK1*, scaffold protein *TBKBP1* has been reported to facilitate the recruitment of *TBK1* to *PKCθ* complexes via *CARD10*, thereby enabling *TBK1* phosphorylation and activation^32^. Activated *TBK1* can, in turn, engage *mTORC1*, promoting *MYC* translation by phosphorylating *4EBP1* and *S6K*, thereby enhancing *MYC* protein output^33,34^. In parallel, TBK1-mediated phosphorylation of *YB-1* has been shown to promote its binding to the *MYC* promoter, leading to increased *MYC* transcription^35^ and providing an additional layer of regulation at the transcriptional level. Previous studies have shown that *TBK1* adaptor proteins can compete for kinase binding, influencing the distribution of *TBK1* activity^36,37^. In parallel, studies of scaffold proteins such as *KSR* suggest that binding to shared components can change signaling by sequestering them^38^. These observations support a general principle in which adaptor-mediated interactions can control the effective availability of a shared kinase across signaling pathways. We therefore propose that a similar competitive sequestration mechanism may operate at the *TBKBP1* locus. In this framework, elevated *TBKBP1* expression may limit the availability of *TBK1* that restrains non-canonical NF-κB signaling, thereby permitting *NIK* stabilization and subsequent activation of *IKKα*, leading to activation of non-canonical NF-κB target genes^39^. In multiple myeloma, aberrant activation of non-canonical NF-κB signaling has been shown to drive epigenomic reprogramming that supports disease progression^40^.

This framework suggests that increased *TBKBP1* expression, driven by germline regulatory variation, may coordinately enhance *MYC*-associated output and non-canonical NF-κB signaling, thereby promoting subclonal expansion and contributing to adverse evolutionary dynamics in multiple myeloma.

## DISCUSSION

Cancer is a biologically unique disease characterized by the cooperative interplay between two distinct genetic compartments: the inherited germline and the acquired somatic genome. While germline variants establish a baseline susceptibility to malignancy, somatic mutations accrued throughout a lifetime serve as the primary drivers of cellular transformation. In this study, we present the first investigation of GSI through a formal evolutionary lens by evaluating associations between germline variants and quantitative descriptors of tumor dynamics. By shifting the analytical focus from mutation-centric endpoints to longitudinal evolutionary processes, our work identifies inherited determinants of tumor fitness and establishes a new dimension for understanding how the germline genome shapes the life history of a cancer. These results were consistent with a recent study of chronic lymphocytic leukemia, reporting associations between sustained subclonal expansion and adverse prognosis^41^.

This study demonstrates that germline variation can influence tumor evolution^5^, extending its role beyond cancer predisposition to the regulation of subclonal dynamics. By linking inherited variation to evolutionary parameters, we identify a layer of genetic control over tumor progression that is enriched in regulatory elements.

Among the loci identified, *TBKBP1* provides a mechanistic example of how germline regulatory variation can shape tumor evolution. Rather than acting through discrete mutational events, variation at this locus appears to influence the pace of subclonal expansion through transcriptional control, linking inherited regulation to evolutionary dynamics and clinical outcome. The convergence of transcriptional, evolutionary, and functional evidence further suggests that germline regulation of signaling pathways, including MYC and NF-κB pathway, may represent a broader mechanism for modulating tumor progression.

Several limitations should be considered. First, although the current sample size provides power to detect loci of moderate effect, it limits resolution at the level of individual variants, necessitating locus-level interpretation. Second, the sequencing depth of the CoMMpass data is relatively shallow for high-confidence germline calling, potentially introducing noise into genotype inference. The mechanistic model proposed here, in which *TBKBP1* modulates *TBK1* availability to influence non-canonical NF-κB signaling, integrates prior biochemical knowledge but remains to be directly tested in myeloma systems. Future studies combining loss-of-function perturbations with biochemical assays will be required to establish causal directionality and to evaluate whether this axis represents a therapeutic vulnerability.

More broadly, this work extends germline–somatic interaction studies from discrete mutational events to the regulation of evolutionary processes. The analytical framework presented here is readily applicable to other cancers and phenotypic outcomes, enabling systematic interrogation of how inherited variation shapes tumor evolution. The identification of loci such as *TBKBP1* and *STAC* further suggests that distinct regulatory mechanisms may converge on the control of subclonal expansion. Together, these findings support the integration of evolutionary parameter inference into germline cancer genomics to uncover heritable determinants of disease progression.

## Funding

This work was supported by National Institutes of Health of the United States R01LM013438.

## METHODS AND MATERIALS

### Cohort and data processing

Germline whole-genome sequencing, Somatic mutation and clinical annotation data were retrieved from the Genomic Data Commons Data Portal^42^. Analyses were restricted to primary tumor samples. Among 895 primary tumors, 869 patients had available demographic and clinical staging information and 506 had both germline and somatic data. Germline variants were quality-controlled using PLINK (--geno 0.05, --maf 0.05, --hwe 1e-6)^43^. Principal components were computed by LD-pruning variants, residualizing per-SNP genotypes against sample source, and performing SVD on the corrected matrix to obtain 10 PCs.

### Evolutionary parameter estimation

Evolutionary parameters were estimated using TEATIME^10^. Estimates were obtained for 418 tumor samples with available sequencing data. Karyotype was obtained from the MMRF CoMMpass dataset. Tumor mutational burden (TMB) was defined as the number of somatic nonsynonymous mutations per megabase of coding sequence, approximated by dividing the total mutation count by 35 Mb^44^. Survival analyses were performed using Cox proportional hazards models, adjusting for age, sex, TMB, and Karyotype. PEER (Probabilistic Estimation of Expression Residuals) factors were computed using the GTEx v8 pipeline^45^. TPM expression values were normalized, and 15 PEER factors were estimated to capture hidden confounders in gene expression. eQTL analysis was performed by regressing each target gene on the genotype adjusted for age, sex, TMB, karyotype, and 15 PEER factors.

### Somatic evolutionary association testing

Per-variant association testing was performed using linear regression of each evolutionary parameter, including mutation rate (*μ*), fitness diversity index (*θ*), subclone selection coefficient (*s*), subclone emergence time (*t*_*f*_) and subclone expansion score (*τ*), on additive, dominant, and recessive genotype coding, adjusted for age, sex, top 10 PCs, TMB, and karyotype. To account for linkage disequilibrium, variants were grouped into LD blocks using a threshold of *R*^2^ ≥ 0.9^46^. Within each block, the minimum P-value (lead SNP) was taken as the block-level test statistic. Genome-wide false discovery rate control was applied to block-level P-values, with blocks at q < 0.05 declared significant^47^.

### Somatic Evolutionary Mediation analysis

To assess whether germline variants influence overall survival through somatic evolutionary parameters, we applied a causal mediation framework. In this framework, the germline variant (exposure *X*) influences an evolutionary parameter (mediator *M*), which in turn may affect the outcome (*Y*). Covariates *C* are included in all models. The mediator model is

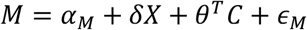

Where *δ* represents the effect of germline variant on the mediator.

For clinical outcomes, overall survival was modelled using a Cox proportional hazards regression:

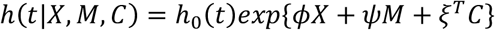

where *ϕ* and *ψ* represents the direct effect of the germline variant and the effect of the mediator, respectively.

Within this framework, the total effect (TE) of X on Y was partitioned into the natural direct effect (NDE) and the natural indirect effect (NIE). The proportion mediated (PM) is defined as NIE/TE. Inference was performed on the log-hazard scale within a causal mediation framework^14,48,49^. To ensure biological interpretability, mediation signals were retained only if (i) the NIE confidence interval excludes zero and (ii) the NIE and TE were directionally concordant. Multiple testing across mediators and variants was controlled using FDR.

### Functional characterization of *TBKBP1*

*TBKBP1* expression across myeloma disease stages was evaluated in two independent GEO microarray cohorts (GSE47552 and GSE6477)^19,20^. MGUS and SMM were combined as a pre-malignant group. Differences in expression between pre-malignant (MGUS/SMM) and multiple myeloma (MM) samples were assessed using the Jonckheere–Terpstra test for ordered trends, with pairwise comparisons performed using Wilcoxon rank-sum tests.

Kaplan–Meier survival analysis was conducted in CoMMpass cohort and an independent GEO cohort (GSE24080)^18^. Analysis was restricted to Total Therapy 3 patients with Serum beta-2 microglobulin level > 5.5 mg/L (ISS stage III). TBKBP1 expression strata were defined as described previously^15,16^.

### Allele-specific expression analysis

Allele-specific expression (ASE) of TBKBP1 was evaluated using RNA-seq data from multiple myeloma cell lines generated at the Mayo Clinic (laboratory of Dr.Leif Bergsagel). Allele-specific read counts were restricted to variants within the TBKBP1 gene body. Only heterozygous sites (genotype 0/1) with ≥5 reads per allele and total depth >10 were retained. Genomic coordinates were lifted from hg19 to hg38 using UCSC LiftOver^50^. Gene-level ASE was quantified using ASEP^21^, which models allelic imbalance across samples using a mixture model with random effects. Each cell line was treated as an independent sample. Statistical significance was assessed by permutation testing (10,000 resamples).

### Cell lines

Human multiple myeloma cell lines KMS11 and MM1.S were obtained from Dr. Leif Bergsagel’s lab. All MM cell lines were maintained in RPMI1640 with 5% fetal bovine serum, 100u/ml penicillin and 100ug/ml streptomycin. Cells tested negative for mycoplasma and cross contamination was ruled out by DNA fingerprinting. All experiments were performed on MM cell lines with less than 20 passages after they were thawed.

### Lenti-vector packaging and infection

Human *TBKBP1* (NM_001394755) was synthesized by Genewiz (115 Corporate Boulevard, South Plainfield, NJ 07080) and cloned into lentivector pCDH-CMV-MCS-EF1-Puro (SBI, Mountain View, CA 94043). Lentivector packaging was described^51^.

Packaging plasmids psPAX2 (Addgene plsmid # 12260) and pMD2.G (Addgene plasmid # 12259) were gifts from Dr. Didier Trono. A modified Graham’s calcium-phosphate method was used for packaging lentiviral vectors^52,53^. Briefly, 293T cells were split into 100mm tissue culture dishes (15 ml total volume medium, no more than 70% confluent before transformation).

Transformation was performed 24 hours later. The 2ml calcium phosphate-DNA co-precipitation solution was made with 100 ul 2.5M CaCl2 and plasmid DNA (18.34 ug for pSPAX2, 7.34 ug for pMD2.G and 24.34 ug for lenti vector), diluted with 1/10 TE, pH 8.0 to the final total volume of 1 ml. Next, one volume of 2xCa/DNA solution was added to one volume of 2x HEPES. After 1 minute, 1.5 ml of the mixed solution was added to the cell culture medium. The next morning the medium was changed with 10 ml fresh DMEM medium and 10% FBS. The viruses were harvested 20 hours after medium change. For infection of MM cell lines, one million cells were added into 6-well plates with 6 ul polybrene (5 mg/ml), 1.5 ml lenti virus and fresh medium to the total volume of 3 ml. Four hours later, 3 ml of fresh medium was added. Next morning the RPMI medium was changed. 72 hours after infection puromycin was added at 3 ug/ml concentration.

### MTT assay

Cellular viability of the MM cell lines was determined by the MTT assay (Sigma-Aldrich, 3050 Spruce St.St. Louis, MO 63103) following the manufacturer’s instructions. Briefly, Seed cells at a concentration of 2 × 103 cells/ well in 100 μl culture medium. Incubate cell cultures for 20 days at 37°C and 5% CO2. Adding 100 ul fresh medium at day 12 and changed medium at day 17.

After the incubation period, add 10 μl of the MTT labeling reagent (final concentration 0.5 mg/ml) to each well. Incubate the microplate for 4 h in a humidified atmosphere (37°C, 5% CO2). Add 100 μl of the Solubilization solution into each well. Allow the plate to stand overnight in the incubator in a humidified atmosphere (37°C, 5% CO2). Check for complete solubilization of the purple formazan crystals and measure the absorbance of the samples using a microplate (ELISA) reader. The wavelength to measure absorbance of the formazan product is 550 nm by the ELISA reader. The reference wavelength is 650 nm. All MTT assay was average of 12 wells on 96 well plate.

### Transcriptomic profiling and pathway enrichment analysis

Paired-end RNA sequencing was performed by Novogene using total RNA from KMS11 and MM1.S cells expressing TBKBP1 or empty vector control. Reads were aligned to the GRCh38 reference genome using STAR^54^, with gene annotation from GENCODE. Gene-level counts were obtained using featureCounts^55^, and genes with fewer than five counts in at least 50% of samples were excluded.

Differential expression analysis was performed in R using DESeq2^56^, incorporating cell line as a covariate in the design formula. Genes with a false discovery rate (FDR) < 0.05 and |log_2_ fold change| > 1 were considered differentially expressed.Gene set enrichment analysis (GSEA), genes with valid Wald statistics and Entrez identifiers were ranked by Wald statistic. Enrichment analysis was conducted against Gene Ontology Biological Process terms using clusterProfiler, with redundant terms collapsed using semantic similarity (Wang method^57^; cutoff = 0.8), and against the MSigDB^58^ Hallmark collection using fgsea^59^.

## Data availability

Only publicly available data were used in this study, and data sources and handling of these data are described above.

## Notes

### Competing Interest Statement

The authors have declared no competing interest.

