## SUPPLEMENTARY_MATERIALS for "Germline regulation of tumor evolutionary dynamics shapes multiple myeloma progression"

**Supplementary Methods**

CoMMpass data processing

Germline whole-genome sequencing (WGS) data from the CoMMpass study were obtained from the Multiple Myeloma Research Foundation CoMMpass project[1]. Adapter trimming was performed using Skewer[2],followed by read pair correction with BBMap[3]. Reads were aligned to the hg19 reference genome using BWA[4], and BAM files were recalibrated with bamUtil[5]. Germline variants were called using Strelka[6]. Genomic coordinates were subsequently lifted over to hg38 using UCSC liftOver[7].

Genomic inflation and QQ plots

To assess the calibration of association statistics, genomic inflation factors (λ) were computed for each evolutionary parameter and genetic model. For each combination, λ was estimated as the ratio of the median observed chi-squared statistic to the expected median under the null distribution. Quantile–quantile plots were generated by comparing observed $-\log_{10}(P)$ values with those expected under a uniform null.

Functional annotation of evolution-informed variants

To evaluate the regulatory potential of germline variants associated with evolutionary parameters, a background set was defined from the full quality-controlled germline variant panel, excluding variants within significant LD blocks. Proximity to transcription start sites was calculated as the distance to the nearest TSS of protein-coding transcripts from GENCODE[8]. Distance distributions between evolution-informed and background variants were compared using a one-sided t-test. Enrichment for expression quantitative trait loci was evaluated by annotating variants for overlap with GTEx v8 whole-blood eQTLs[9], with proportional differences assessed using chi-squared tests.

Survival stratification by gene expression

Expression strata were defined by scanning candidate cut-points across the 20th–80th percentile range of $log_2$(TPM + 1) values and selecting the threshold that minimized the log-rank P value [10,11].

cis-eQTL analysis and locus visualisation

To assess whether germline variants at candidate loci regulate gene expression in multiple myeloma, cis-eQTL analyses were performed within a ±500 kb window of each gene locus across cohorts. In the CoMMpass cohort, germline SNP genotypes were integrated with matched gene expression while adjusting for age, gender, TMB and Karyotype. Associations between SNP genotype and gene expression were tested using linear regression, and SNPs with P<0.05 were considered nominally significant. Effect sizes were normalized. Linkage disequilibrium ($R^2$) among significant variants was estimated using the NIH LD matrix[12] with a European reference population.

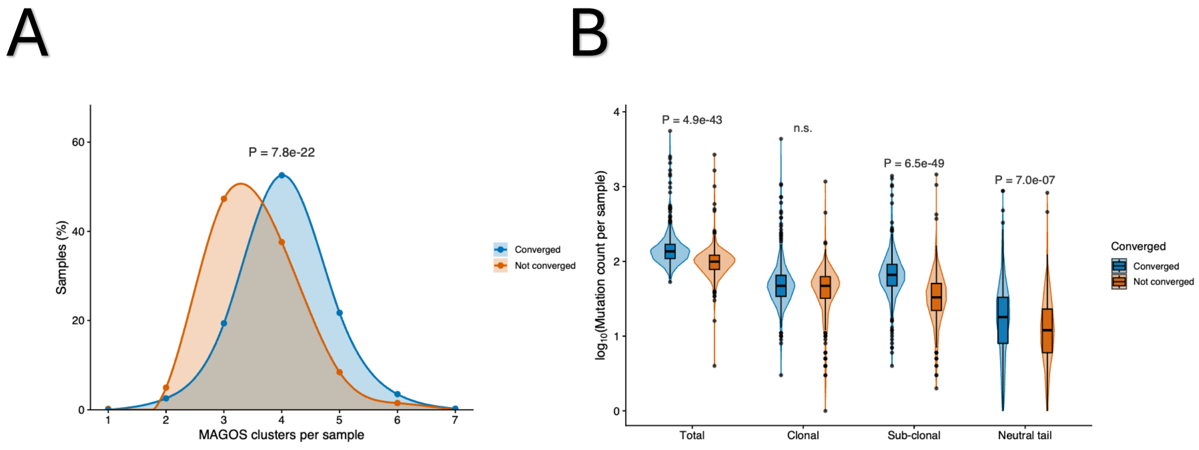


**Supplementary Figure 1.** TEATIME convergence is associated with mutation cluster structure and cluster-level mutation burden. A. Distribution of the number of mutation clusters in convergent and non-convergent TEATIME cases. B. Distribution of mutation counts within inferred clusters in convergent and non-convergent TEATIME cases.

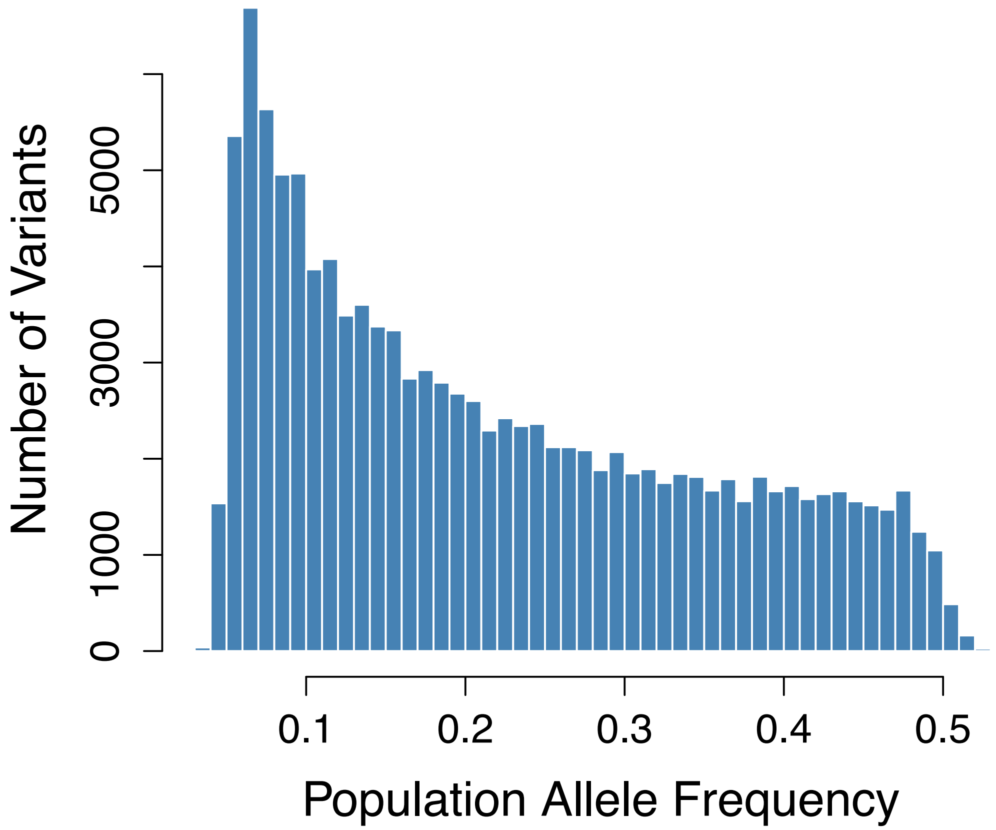


**Supplementary Figure 2.** Population allele Frequency in TEATIME converged Samples.

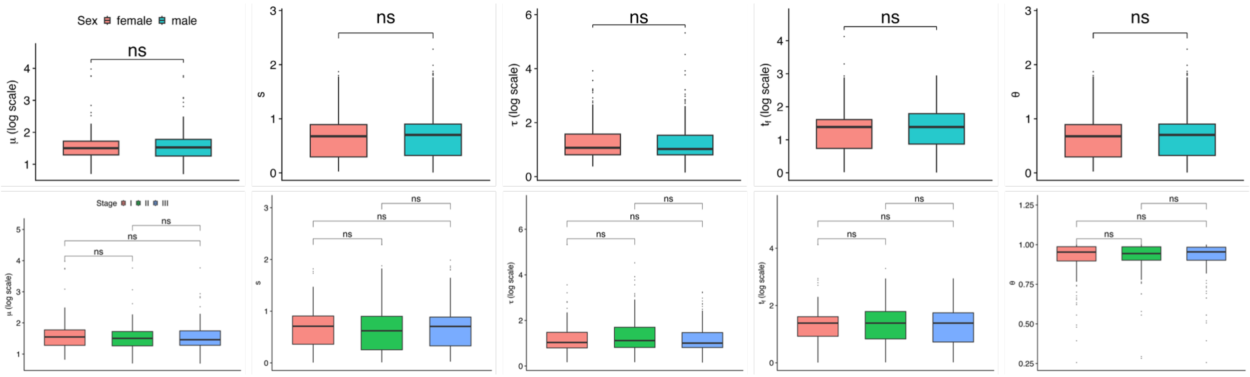


**Supplementary Figure 3.** Stratified analyses of evolutionary parameters by sex and disease stage.

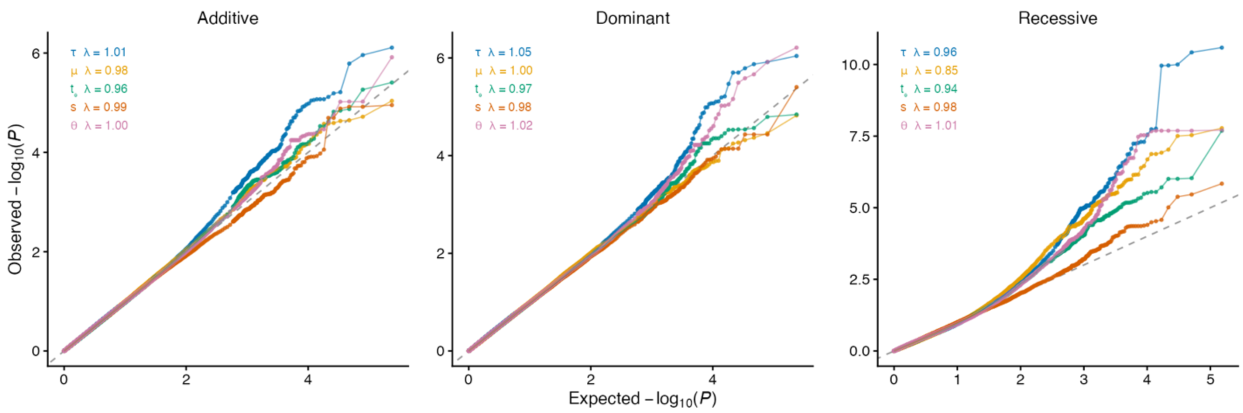


**Supplementary Fig 4.** Quantile–quantile analysis of three models.

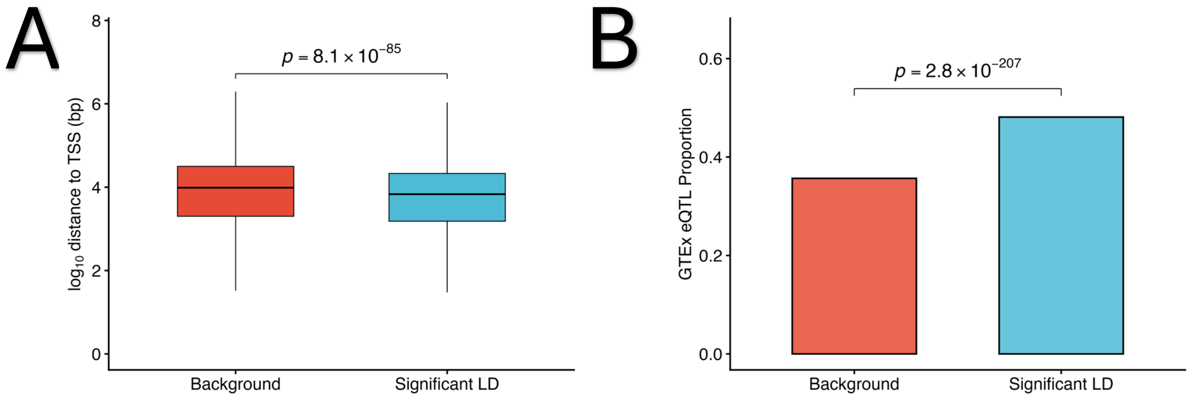


**Supplementary Fig 5.** Regulatory enrichment of germline loci. (**A**) Distribution of transcription start site (TSS) distances across parameters (**B**) Proportion of loci that overlap with GTEx whole blood eQTLs across parameters

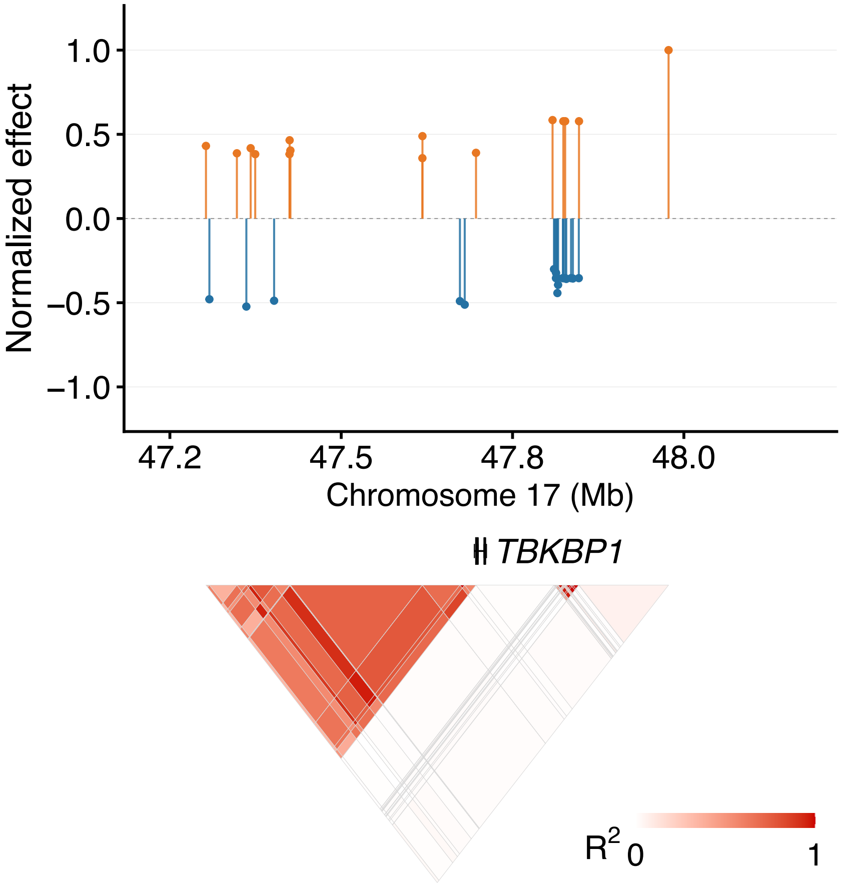


**Supplementary Fig 6.** The top panel shows normalized effect sizes of eQTLs. Gene models are shown in the middle panels. The bottom panel shows local LD structure ($R^2$).

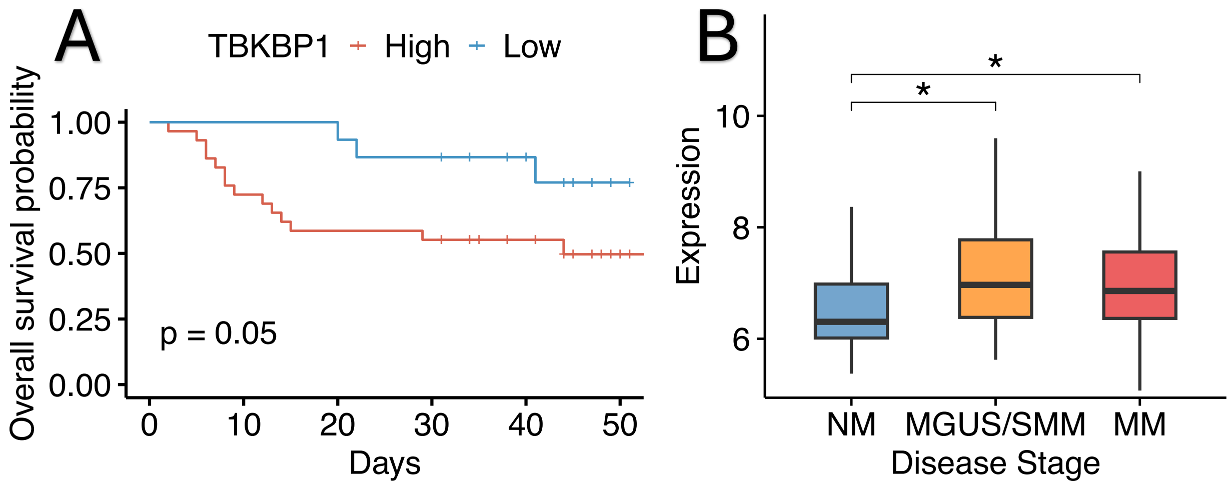


**Supplementary Fig 7.** Independent cohort validation. (A) GSE24080 patients Kaplan–Meier overall survival curves (B) GSE6477 *TBKBP1* expression in different disease stage.

**Supplementary Table 1** LD blocks significantly associated with tumor evolutionary parameters

| phenotype | moc | locus | lead_variant | qvalue | beta | phenotyp | moc | locus.2 | lead_variant.2 | qvalue.2 | beta.2 | phenotyp | mo | locus.3 | lead_variant.3 | qvalue.3 | beta.3 | phenoty | mo | locus.4 | lead_variant.4 | qvalue.4 | beta.4 | phenotyp | mc | locus.5 | lead_variant.5 | qvalue.5 | beta.5 | phenotyp | moc | locus.6 | lead_variant.6 | qvalue.6 | beta.6 | phenoty | mc | locus.7 | lead_variant.7 | qvalue.7 | beta.7 |
|---|---|---|---|---|---|---|---|---|---|---|---|---|---|---|---|---|---|---|---|---|---|---|---|---|---|---|---|---|---|---|---|---|---|---|---|---|---|---|---|---|---|
| entropy | R | 13q31.2 | chr13:89,360,933 | 0 | -0.422 | mu | R | 19q13.43 | chr19:57,633,151 | 0.0072 | 5.711 | tau | R | 19q13.42 | chr19:54,148,468 | 0.0187 | 7.4626 | tf | R | 15q24.2 | chr15:75,579,180 | 0.0275 | 5.8182 | entropy | R | 3p21.31 | chr3:48,463,903 | 0.0331 | -0.111 | tf | D | 20p12.2 | chr20:10,407,858 | 0.0385 | 2.8035 | tf | A | 2q33.1 | chr2:201,218,018 | 0.045 | -2.187 |
| entropy | R | 9q34.3 | chr9:136,740,043 | 0 | -0.412 | mu | R | 1q24.1 | chr1:166,989,700 | 0.0072 | 4.232 | entropy | R | 22q12.3 | chr22:32,193,103 | 0.0187 | -0.281 | tf | R | 16p12.1 | chr16:24,827,667 | 0.0275 | 3.4186 | tf | R | 5q11.2 | chr5:55,137,565 | 0.0331 | 2.5183 | tf | D | 4q32.3 | chr4:165,484,648 | 0.0385 | 2.3674 | tf | A | 5q14.3 | chr5:83,538,811 | 0.045 | -1.159 |
| entropy | R | 16p11.2 | chr16:28,981,728 | 1e-04 | -0.343 | tau | R | 1p31.1 | chr1:78,537,838 | 0.0072 | 8.016 | mu | R | 22q13.2 | chr22:43,218,217 | 0.0187 | 5.7638 | tau | R | 17p13.2 | chr17:5,421,268 | 0.0275 | 6.7472 | mu | R | 11q21 | chr11:94,464,402 | 0.0332 | 2.1398 | tf | R | 12q24.11 | chr12:109,445,449 | 0.0388 | 2.6682 | tf | A | 11q22.1 | chr11:101,209,684 | 0.0452 | 2.5014 |
| tau | R | 18q23 | chr18:80,138,359 | 1e-04 | 11.83 | mu | R | 12q24.33 | chr12:132,675,970 | 0.0074 | 2.558 | entropy | R | 2p13.3 | chr2:70,936,043 | 0.0187 | -0.198 | tf | R | 19q13.33 | chr19:49,880,379 | 0.0275 | -2.0391 | tf | R | 19p13.2 | chr19:12,272,275 | 0.0332 | 5.4657 | tf | R | 14q24.3 | chr14:77,508,760 | 0.0388 | 3.5084 | tau | R | 14q11.2 | chr14:22,834,885 | 0.0453 | 5.3579 |
| tau | R | 19q13.12 | chr19:35,142,599 | 1e-04 | 7.285 | mu | R | 6p21.33 | chr6:31,279,192 | 0.0074 | 5.489 | entropy | R | 9q34.3 | chr9:137,435,166 | 0.0187 | -0.172 | tau | R | 19q13.43 | chr19:56,205,984 | 0.0275 | 3.5381 | tau | A | 10q24.2 | chr10:99,612,654 | 0.0336 | 2.8585 | tf | A | 11p13 | chr11:34,492,013 | 0.0391 | 2.3916 | tf | R | 11q13.4 | chr11:72,579,585 | 0.0456 | 3.4421 |
| entropy | R | 19p13.2 | chr19:6,981,752 | 1e-04 | -0.403 | tau | R | 10q24.1 | chr10:95,637,470 | 0.0081 | 4.875 | entropy | R | 12q24.13 | chr12:113,169,69 | 0.0189 | -0.197 | tf | R | 5p13.1 | chr5:41,586,109 | 0.0275 | 4.0725 | tau | A | 10q23.31 | chr10:88,778,185 | 0.0336 | 4.9018 | mu | R | 15q25.3 | chr15:85,710,537 | 0.0391 | 4.6352 | tf | R | 2q21.3 | chr2:134,987,975 | 0.0456 | 2.583 |
| tau | R | 1q23.3 | chr1:162,381,664 | 1e-04 | 14.22 | entropy | R | 20p13 | chr20:2,572,238 | 0.0081 | -0.3 | tf | R | 19p13.12 | chr19:15,897,289 | 0.0189 | 3.8054 | tf | R | 6q25.3 | chr6:158,626,350 | 0.0275 | 3.676 | tau | A | 11q24.2 | chr11:124,185,670 | 0.0336 | 2.1135 | tau | R | 19q13.33 | chr19:50,244,171 | 0.0393 | 5.1265 | tf | R | 14q23.1 | chr14:59,610,065 | 0.0457 | 3.2048 |
| entropy | R | 20q11.21 | chr20:31,831,248 | 1e-04 | -0.385 | tf | R | 22q12.2 | chr22:29,696,594 | 0.0081 | 5.749 | tf | R | 2q37.1 | chr2:231,048,680 | 0.0189 | 5.1161 | tau | R | 17q25.1 | chr17:73,201,300 | 0.0276 | 8.9166 | tau | A | 12q23.3 | chr12:103,793,641 | 0.0336 | 5.1768 | tau | R | 19q13.42 | chr19:54,451,197 | 0.0393 | 5.7191 | tau | D | 1q21.3 | chr1:151,001,835 | 0.0457 | 5.3519 |
| tau | R | 2q11.2 | chr2:102,016,112 | 1e-04 | 14.05 | tau | D | 19p13.3 | chr19:3,633,117 | 0.0083 | 4.792 | tf | R | 11p11.2 | chr11:48,217,328 | 0.0191 | 5.56 | tf | D | 2q33.1 | chr2:197,081,668 | 0.0276 | 3.7664 | tau | A | 12q23.3 | chr12:106,844,099 | 0.0336 | 4.5493 | tau | R | 21q22.3 | chr21:42,485,777 | 0.0393 | 4.8748 | tau | A | 11p15.1 | chr11:20,607,618 | 0.0459 | 4.3154 |
| entropy | R | 4q32.1 | chr4:155,785,260 | 1e-04 | -0.347 | mu | R | 2q23.3 | chr2:150,468,789 | 0.0084 | 6.841 | tf | R | 16p13.3 | chr16:7,051,975 | 0.0191 | 6.1691 | tf | R | 5q22.3 | chr5:115,523,909 | 0.0276 | 5.0364 | tau | A | 12p13.2 | chr12:12,094,682 | 0.0336 | 2.3888 | entropy | R | 11p11.12 | chr11:49,820,165 | 0.0395 | -0.24 | tau | A | 15q13.3 | chr15:33,154,678 | 0.0459 | -2.267 |
| mu | R | 7q36.1 | chr7:150,743,626 | 1e-04 | 8.701 | mu | R | 4q32.3 | chr4:168,928,086 | 0.0084 | 6.483 | tf | R | 19p13.11 | chr19:17,226,924 | 0.0191 | 5.095 | mu | R | 14q11.2 | chr14:23,099,817 | 0.0279 | -7.4674 | tau | A | 12q12 | chr12:43,453,880 | 0.0336 | 4.7024 | entropy | R | 13q12.12 | chr13:24,868,783 | 0.0395 | -0.113 | mu | R | 1q23.1 | chr1:156,669,961 | 0.0459 | -1.876 |
| entropy | R | 8q22.1 | chr8:93,703,737 | 1e-04 | -0.394 | mu | R | 9q22.32 | chr9:95,928,855 | 0.0084 | 5.838 | tf | R | 22q12.3 | chr22:36,506,699 | 0.0191 | 6.2912 | tf | D | 20p13 | chr20:275,932 | 0.0279 | -2.4597 | tau | A | 12q13.12 | chr12:50,885,584 | 0.0336 | 3.4976 | entropy | R | 18p11.22 | chr18:10,471,735 | 0.0395 | -0.249 | entropy | R | 1p36.33 | chr1:1,452,346 | 0.0461 | -0.229 |
| tau | R | 9q34.2 | chr9:133,520,331 | 1e-04 | 11.39 | mu | R | 9q22.32 | chr9:95,972,609 | 0.0084 | 5.838 | tf | R | 2p25.1 | chr2:10,784,398 | 0.0191 | 2.3718 | mu | R | 20q11.23 | chr20:35,855,251 | 0.0279 | 3.7589 | tau | A | 12q13.13 | chr12:53,062,096 | 0.0336 | 5.1193 | entropy | R | 20q11.21 | chr20:32,997,626 | 0.0395 | -0.245 | entropy | R | 3q13.13 | chr3:108,757,127 | 0.0461 | -0.131 |
| entropy | R | 14q13.1 | chr14:33,213,919 | 2e-04 | -0.323 | mu | R | 11q12.1 | chr11:56,232,995 | 0.0085 | 4.716 | entropy | R | 10q26.2 | chr10:125,795,004 | 0.0195 | -0.117 | tf | R | 11q24.3 | chr11:128,902,55 | 0.0281 | 4.0754 | tau | A | 13q34 | chr13:113,768,927 | 0.0336 | 2.3879 | tf | A | 4q34.1 | chr4:173,616,205 | 0.0395 | -1.262 | entropy | R | 6p25.1 | chr6:4,715,648 | 0.0461 | -0.11 |
| mu | R | 19p13.2 | chr19:7,900,454 | 2e-04 | 10.93 | mu | R | 17p12 | chr17:14,770,457 | 0.0085 | 2.919 | entropy | R | 11q24.3 | chr11:128,522,042 | 0.0195 | -0.287 | tf | R | 21q22.3 | chr21:46,194,555 | 0.0281 | 3.5647 | tau | A | 13q14.13 | chr13:45,702,368 | 0.0336 | 2.7742 | entropy | R | 6q27 | chr6:168,294,629 | 0.0395 | -0.097 | s | D | 10q21.1 | chr10:51,695,802 | 0.0462 | 0.4787 |
| mu | R | 20q11.22 | chr20:35,543,647 | 2e-04 | 8.604 | mu | R | 19q13.2 | chr19:39,416,396 | 0.0085 | 4.948 | entropy | R | 12q24.31 | chr12:122,208,273 | 0.0195 | -0.127 | tf | R | 14q32.2 | chr14:96,534,003 | 0.0282 | 4.0512 | tau | A | 15q26.3 | chr15:100,063,814 | 0.0336 | 4.0964 | tf | R | 12q24.21 | chr12:114,670,331 | 0.0399 | 2.9428 | s | D | 11q23.3 | chr11:118,307,292 | 0.0462 | -0.308 |
| entropy | R | 6q22.1 | chr6:117,404,285 | 2e-04 | -0.374 | mu | R | 19q13.2 | chr19:39,518,826 | 0.0085 | 6.675 | entropy | R | 16p12.1 | chr16:27,225,082 | 0.0195 | -0.121 | tf | R | 18q12.2 | chr18:35,974,812 | 0.0282 | 3.9725 | tau | A | 16q22.1 | chr16:70,514,316 | 0.0336 | 4.3599 | tf | R | 12q24.33 | chr12:133,041,405 | 0.0399 | 3.9585 | s | D | 11p15.4 | chr11:7,995,183 | 0.0462 | -0.292 |
| entropy | R | 16p11.2 | chr16:28,933,075 | 4e-04 | -0.262 | mu | R | 19q13.2 | chr19:39,539,030 | 0.0085 | 6.675 | entropy | R | 20q12 | chr20:41,362,060 | 0.0195 | -0.252 | tf | R | 1q21.3 | chr1:154,378,113 | 0.0282 | 4.5891 | tau | A | 18q11.2 | chr18:23,845,027 | 0.0336 | 4.809 | tf | R | 6p21.33 | chr6:30,723,161 | 0.0399 | 5.3075 | s | D | 13q14.11 | chr13:42,302,154 | 0.0462 | 0.3582 |
| tau | R | 1p34.1 | chr1:45,510,475 | 4e-04 | 6.96 | mu | R | 1p36.13 | chr1:19,257,142 | 0.0085 | 5.32 | tau | D | 11q24.2 | chr11:124,225,542 | 0.0196 | -3.636 | tf | R | 2q37.3 | chr2:241,246,976 | 0.0282 | 5.1729 | tau | A | 1p13.2 | chr1:112,701,811 | 0.0336 | 2.9328 | mu | R | 2p21 | chr2:47,006,610 | 0.0401 | 4.6639 | s | D | 16q23.3 | chr16:81,869,385 | 0.0462 | 0.2831 |
| tau | R | 6p21.1 | chr6:43,672,302 | 4e-04 | 11.44 | mu | R | 5q15 | chr5:96,757,737 | 0.0085 | 5.458 | mu | R | 16q24.3 | chr16:88,835,775 | 0.0196 | 3.5709 | tf | R | 3q25.1 | chr3:151,824,949 | 0.0282 | 3.8628 | tau | A | 1q21.3 | chr1:151,001,835 | 0.0336 | 5.271 | mu | R | 4q21.21 | chr4:78,364,200 | 0.0401 | 5.1684 | s | D | 4q21.21 | chr4:81,053,655 | 0.0462 | 0.3599 |
| tau | R | 7p13 | chr7:44,050,305 | 4e-04 | 10.69 | mu | R | 6p21.32 | chr6:32,850,997 | 0.0085 | 6.238 | tau | R | 17p13.2 | chr17:5,443,279 | 0.0196 | 8.3062 | tf | R | 4p16.1 | chr4:7,689,692 | 0.0282 | 2.446 | tau | A | 1q25.1 | chr1:175,014,497 | 0.0336 | 2.2712 | mu | R | 5p15.33 | chr5:251,592 | 0.0401 | 4.3657 | s | D | 5p15.33 | chr5:482,538 | 0.0462 | 0.3374 |
| entropy | R | 9p13.3 | chr9:33,662,391 | 5e-04 | -0.215 | mu | R | 6p21.32 | chr6:33,085,918 | 0.0085 | 6.611 | mu | R | 3q29 | chr3:196,935,404 | 0.0197 | 4.9701 | tf | R | 6q14.1 | chr6:75,921,758 | 0.0282 | 4.6548 | tau | A | 21q22.12 | chr21:36,048,352 | 0.0336 | 4.0237 | mu | R | 8q24.13 | chr8:124,143,465 | 0.0401 | -3.688 | s | D | 8p21.3 | chr8:19,395,091 | 0.0462 | -0.326 |
| mu | R | 11p11.2 | chr11:43,886,680 | 7e-04 | 7.936 | mu | R | 7q36.1 | chr7:150,749,604 | 0.0085 | 4.481 | entropy | R | 15q22.2 | chr15:60,421,457 | 0.0198 | -0.133 | mu | R | 14q32.13 | chr14:94,309,884 | 0.0283 | -2.0902 | tau | A | 2p11.2 | chr2:85,616,308 | 0.0336 | 4.7178 | mu | R | 8q24.23 | chr8:138,141,131 | 0.0401 | 2.481 | tf | R | 9q33.1 | chr9:116,401,230 | 0.0464 | 2.8002 |
| tau | R | 3q12.2 | chr3:100,364,013 | 7e-04 | 7.572 | mu | R | 1p13.1 | chr1:116,989,750 | 0.0088 | 3.765 | entropy | R | 20p11.23 | chr20:18,567,781 | 0.0198 | -0.134 | tf | R | 2p23.2 | chr2:28,948,219 | 0.0283 | 4.3215 | tau | A | 3q13.13 | chr3:108,376,888 | 0.0336 | 4.3805 | tf | R | 10q24.33 | chr10:103,855,316 | 0.0402 | 2.9159 | tau | A | 1p34.1 | chr1:45,620,657 | 0.0466 | 2.2765 |
| tau | R | 10q25.3 | chr10:113,774,441 | 8e-04 | 12.36 | entropy | R | 7q22.1 | chr7:102,273,527 | 0.0088 | -0.298 | entropy | R | 21q22.3 | chr21:46,538,744 | 0.0198 | -0.212 | mu | R | 3q13.2 | chr3:113,274,520 | 0.0283 | 4.781 | tau | A | 5q33.1 | chr5:151,492,234 | 0.0336 | 2.6628 | entropy | R | 11p14.2 | chr11:26,565,254 | 0.0402 | -0.118 | tau | R | 7q36.1 | chr7:148,417,773 | 0.0466 | 3.2337 |
| tau | R | 12p13.2 | chr12:12,094,660 | 8e-04 | 7.397 | entropy | R | 9p13.3 | chr9:33,572,231 | 0.0088 | -0.247 | entropy | R | 4q35.1 | chr4:185,161,245 | 0.0198 | -0.109 | tf | R | 11q24.3 | chr11:130,877,49 | 0.0284 | 1.7189 | tau | A | 5q35.3 | chr5:177,600,404 | 0.0336 | 4.6359 | tau | A | 12q23.3 | chr12:108,538,003 | 0.0402 | 2.766 | entropy | R | 5q34 | chr5:162,142,424 | 0.0467 | -0.241 |
| tau | R | 15q21.3 | chr15:57,550,243 | 8e-04 | 10.75 | tau | R | 1q25.2 | chr1:179,842,369 | 0.0089 | 10.17 | mu | R | 14q32.33 | chr14:105,986,730 | 0.0199 | 4.2853 | entropy | R | 15q21.1 | chr15:45,597,920 | 0.0285 | -0.2522 | tau | A | 6q23.2 | chr6:134,259,980 | 0.0336 | 2.3625 | tau | R | 12q24.31 | chr12:122,716,744 | 0.0402 | 4.5345 | tf | A | 2p13.3 | chr2:70,259,275 | 0.0468 | 1.8467 |
| mu | R | 5q33.1 | chr5:151,065,282 | 9e-04 | 7.757 | tau | R | 1p36.11 | chr1:26,470,431 | 0.0089 | 5.809 | tau | R | 3p22.3 | chr3:36,380,643 | 0.0199 | 8.4155 | entropy | A | 15q25.3 | chr15:85,717,897 | 0.0285 | -0.1354 | tau | A | 6q24.2 | chr6:142,220,632 | 0.0336 | 4.0655 | tau | A | 12q24.33 | chr12:130,872,027 | 0.0402 | 3.9363 | tf | D | 22q13.31 | chr22:44,735,173 | 0.0469 | 2.9351 |
| tau | R | 13q34 | chr13:112,882,153 | 0.001 | 9.506 | entropy | R | 20p13 | chr20:1,630,144 | 0.0091 | -0.159 | mu | R | 7q22.1 | chr7:100,467,096 | 0.0199 | 2.2128 | entropy | A | 6p21.33 | chr6:31,356,368 | 0.0285 | -0.0933 | tau | A | 6p25.3 | chr6:549,390 | 0.0336 | -2.145 | entropy | R | 12q13.12 | chr12:49,958,776 | 0.0402 | -0.212 | tau | A | 6q22.1 | chr6:116,452,402 | 0.0469 | 2.7969 |
| tau | R | 1p13.2 | chr1:112,699,866 | 0.001 | 6.909 | mu | R | 20q11.22 | chr20:35,579,460 | 0.0092 | 4.942 | tf | R | 22q13.31 | chr22:44,859,750 | 0.0201 | 6.1093 | tf | R | 22q13.31 | chr22:45,922,405 | 0.0286 | 4.916 | tau | A | 7q36.1 | chr7:150,742,955 | 0.0336 | 2.2009 | tau | A | 12p13.31 | chr12:5,985,618 | 0.0402 | 2.5835 | mu | R | 11p15.5 | chr11:609,068 | 0.047 | 3.3441 |
| tau | R | 2q33.2 | chr2:203,867,991 | 0.001 | 6.163 | tau | A | 12q23.3 | chr12:107,635,409 | 0.0093 | 4.01 | s | R | 15q25.3 | chr15:85,692,004 | 0.0202 | 0.9577 | mu | R | 12q14.3 | chr12:66,420,707 | 0.0287 | -2.1018 | tau | A | 7p13 | chr7:44,050,305 | 0.0336 | 2.6355 | tau | R | 13q34 | chr13:114,013,085 | 0.0402 | 5.0289 | mu | R | 14q24.3 | chr14:76,810,279 | 0.047 | 5.3137 |
| tau | R | 7p22.3 | chr7:2,355,356 | 0.001 | 8.664 | tau | A | 16p13.3 | chr16:2,317,335 | 0.0093 | 4.323 | tf | R | 18q21.33 | chr18:63,603,831 | 0.0202 | 4.4754 | mu | R | 1p36.13 | chr1:19,193,288 | 0.0287 | 2.1837 | tau | A | 8q12.3 | chr8:61,714,445 | 0.0336 | 2.4801 | tau | A | 17p12 | chr17:14,768,804 | 0.0402 | 3.723 | mu | R | 17q25.1 | chr17:75,891,915 | 0.047 | 4.3976 |
| mu | R | 20q11.21 | chr20:33,002,939 | 0.0011 | 7.712 | tau | A | 17p12 | chr17:14,077,333 | 0.0093 | 5.264 | tf | R | 1q25.2 | chr1:179,483,140 | 0.0202 | 6.1671 | s | R | 2q37.3 | chr2:241,114,988 | 0.0288 | 1.135 | entropy | R | 9q31.2 | chr9:105,461,505 | 0.0336 | -0.255 | tau | R | 17p13.2 | chr17:4,639,422 | 0.0402 | 5.2957 | mu | R | 4q32.3 | chr4:168,177,763 | 0.047 | 3.8857 |
| mu | R | 20q11.22 | chr20:35,502,691 | 0.0011 | 8.444 | tau | R | 2q37.3 | chr2:239,578,479 | 0.0094 | 4.092 | s | R | 1q44 | chr1:246,855,975 | 0.0202 | 0.7456 | tf | R | 10p11.23 | chr10:29,533,230 | 0.0289 | 2.3586 | tau | A | 9q34.2 | chr9:133,520,315 | 0.0336 | 2.5738 | tau | R | 19q13.12 | chr19:35,125,759 | 0.0402 | 4.3452 | tf | R | 11p15.4 | chr11:8,683,933 | 0.0474 | 5.2263 |
| tau | R | 19p13.12 | chr19:15,897,289 | 0.0012 | 8.523 | tf | R | 6q16.2 | chr6:99,568,924 | 0.0094 | 6.142 | s | R | 20q13.33 | chr20:62,394,395 | 0.0202 | 0.8673 | tf | R | 12q24.31 | chr12:121,761,74 | 0.0289 | 5.9022 | tf | A | 22q12.2 | chr22:31,611,167 | 0.0337 | 2.6746 | tf | R | 19q13.32 | chr19:45,823,795 | 0.0402 | 5.3951 | tf | A | 17p12 | chr17:15,230,089 | 0.0474 | 2.2381 |
| tau | R | 8q12.3 | chr8:61,714,445 | 0.0012 | 8.523 | mu | R | 3q21.3 | chr3:128,861,742 | 0.0095 | 3.406 | s | R | 22q11.21 | chr22:19,132,238 | 0.0202 | 0.7173 | tf | R | 15q26.3 | chr15:101,654,08 | 0.0291 | 3.7734 | tf | A | 3p25.3 | chr3:10,002,578 | 0.0339 | -1.129 | tau | R | 19q13.41 | chr19:51,627,235 | 0.0402 | 8.6198 | tf | R | 17p11.2 | chr17:18,930,945 | 0.0474 | 4.1769 |
| mu | R | 11q13.2 | chr11:68,030,896 | 0.0014 | 6.873 | mu | R | 4p15.31 | chr4:20,713,621 | 0.0095 | 5.437 | tf | R | 5q11.2 | chr5:53,072,642 | 0.0202 | 6.1398 | tf | A | 10p12.1 | chr10:27,208,262 | 0.0293 | -2.7911 | tf | A | 5q35.1 | chr5:169,594,122 | 0.034 | 2.4081 | tau | A | 1q32.2 | chr1:207,474,828 | 0.0402 | 3.6259 | tau | R | 14q12 | chr14:24,417,895 | 0.0475 | 7.3235 |
| tau | R | 14q11.2 | chr14:22,169,058 | 0.0014 | 9.899 | entropy | R | 10q26.2 | chr10:125,836,043 | 0.0099 | -0.117 | s | R | 6p21.32 | chr6:32,288,696 | 0.0202 | 0.6992 | tf | A | 11p15.4 | chr11:5,403,302 | 0.0293 | 2.0801 | mu | R | 19p13.2 | chr19:7,616,465 | 0.0343 | 3.979 | tau | R | 1q41 | chr1:220,317,470 | 0.0402 | 3.3791 | tau | R | 19p12 | chr19:21,328,982 | 0.0475 | 5.8613 |
| tau | R | 17p11.2 | chr17:16,939,677 | 0.0014 | 12.21 | tau | R | 19p13.11 | chr19:18,177,512 | 0.0099 | 7.917 | tf | R | 6p25.3 | chr6:410,493 | 0.0202 | 4.5248 | tf | A | 12q23.3 | chr12:103,982,91 | 0.0293 | 3.169 | tf | R | 14q32.12 | chr14:92,685,839 | 0.0345 | 2.0703 | tau | A | 1p34.1 | chr1:44,841,834 | 0.0402 | 2.045 | tau | R | 22q11.23 | chr22:23,927,818 | 0.0475 | 5.5424 |
| tau | R | 19q13.42 | chr19:53,407,659 | 0.0014 | 10.09 | tau | A | 1p34.1 | chr1:45,332,883 | 0.0099 | 4.622 | entropy | R | 10q23.31 | chr10:89,402,920 | 0.0203 | -0.232 | tf | A | 14q13.2 | chr14:34,939,442 | 0.0293 | 1.5549 | mu | R | 10q23.31 | chr10:88,673,588 | 0.0346 | 5.0546 | tau | A | 1p36.23 | chr1:7,781,270 | 0.0402 | 3.5559 | tau | R | 2q13 | chr2:110,184,919 | 0.0475 | 4.357 |
| entropy | R | 22q11.21 | chr22:19,964,978 | 0.0014 | -0.199 | entropy | R | 3q27.2 | chr3:185,632,918 | 0.0099 | -0.157 | entropy | R | 12q23.3 | chr12:108,542,708 | 0.0203 | -0.294 | tf | A | 15q26.3 | chr15:99,137,940 | 0.0293 | 2.0862 | mu | R | 11q13.2 | chr11:68,018,929 | 0.0346 | 3.7968 | tau | R | 2q37.3 | chr2:238,098,922 | 0.0402 | 5.7197 | tau | R | 2q33.1 | chr2:202,201,325 | 0.0475 | 4.5361 |
| mu | R | 22q11.23 | chr22:23,743,060 | 0.0014 | 6.42 | mu | R | 7p12.3 | chr7:45,714,983 | 0.0099 | 5.376 | entropy | R | 12q24.13 | chr12:113,277,446 | 0.0203 | -0.168 | tf | A | 17q21.2 | chr17:41,515,472 | 0.0293 | 1.6642 | mu | R | 16p11.2 | chr16:28,605,736 | 0.0346 | 2.7141 | tau | R | 2p25.1 | chr2:9,443,793 | 0.0402 | 3.7926 | tau | A | 3p22.1 | chr3:42,645,786 | 0.0475 | 2.0647 |
| mu | R | 2p22.2 | chr2:37,179,501 | 0.0014 | 5.862 | entropy | R | 10q24.32 | chr10:102,424,456 | 0.0101 | -0.226 | s | R | 2p12 | chr2:79,028,462 | 0.0203 | -0.355 | tf | A | 19p13.2 | chr19:12,440,915 | 0.0293 | 3.0704 | mu | R | 16p11.2 | chr16:28,904,206 | 0.0346 | 3.5056 | entropy | R | 3q13.33 | chr3:119,643,522 | 0.0402 | -0.136 | tau | R | 7q22.1 | chr7:100,417,223 | 0.0475 | 7.2169 |
| mu | R | 4q26 | chr4:119,060,823 | 0.0014 | 7.623 | entropy | R | 18q21.32 | chr18:59,296,535 | 0.0101 | -0.287 | entropy | R | 3q25.33 | chr3:160,436,839 | 0.0203 | -0.268 | tf | A | 1q24.1 | chr1:166,852,946 | 0.0293 | -1.5375 | mu | R | 2p13.2 | chr2:71,398,637 | 0.0346 | -4.493 | entropy | R | 4q35.2 | chr4:188,097,332 | 0.0402 | -0.253 | tau | R | 7p15.2 | chr7:27,525,587 | 0.0475 | 5.114 |
| tau | R | 13q34 | chr13:113,804,775 | 0.0015 | 11.62 | entropy | R | 1q32.1 | chr1:205,062,616 | 0.0101 | -0.286 | entropy | R | 6q25.3 | chr6:158,237,830 | 0.0203 | -0.283 | entropy | D | 1q32.1 | chr1:207,098,927 | 0.0293 | -0.1247 | mu | R | 3p22.2 | chr3:38,084,100 | 0.0346 | 4.7326 | entropy | R | 5p15.2 | chr5:10,680,435 | 0.0402 | -0.157 | tau | R | 7p22.1 | chr7:5,232,546 | 0.0475 | 8.6969 |
| mu | R | 15q21.2 | chr15:52,128,294 | 0.0015 | 7.726 | entropy | R | 1q32.1 | chr1:205,070,167 | 0.0101 | -0.286 | entropy | R | 9q31.1 | chr9:101,421,740 | 0.0203 | -0.106 | tf | A | 22q13.31 | chr22:44,735,173 | 0.0293 | 2.8713 | tau | R | 6p22.1 | chr6:29,945,489 | 0.0346 | 6.6299 | entropy | R | 6q22.1 | chr6:116,809,381 | 0.0402 | -0.121 | tau | R | 7q22.1 | chr7:100,319,793 | 0.0475 | 7.2169 |
| tau | R | 1q23.2 | chr1:159,806,576 | 0.0015 | 11.78 | entropy | R | 1p32.3 | chr1:53,594,343 | 0.0101 | -0.176 | entropy | R | 14q12 | chr14:24,330,081 | 0.0204 | -0.216 | tf | A | 2q21.3 | chr2:134,451,250 | 0.0293 | 2.7963 | entropy | R | 2q35 | chr2:216,446,886 | 0.0347 | -0.261 | tau | A | 7q22.1 | chr7:100,367,166 | 0.0402 | 2.6518 | tf | A | 10p11.23 | chr10:29,533,230 | 0.0476 | 1.1792 |
| tau | R | 1p31.3 | chr1:62,039,169 | 0.0015 | 8.132 | entropy | R | 3q27.2 | chr3:185,629,671 | 0.0101 | -0.156 | entropy | R | 6q25.3 | chr6:158,767,096 | 0.0204 | -0.14 | tf | A | 2p24.1 | chr2:20,239,735 | 0.0293 | 3.1503 | tau | A | 12q24.11 | chr12:109,073,451 | 0.0348 | 2.537 | tau | R | 8q23.1 | chr8:108,248,547 | 0.0402 | 3.9957 | tf | A | 10q22.3 | chr10:79,920,193 | 0.0476 | 1.6313 |
| mu | R | 20q11.22 | chr20:35,632,065 | 0.0015 | 6.1 | entropy | R | 4q35.2 | chr4:188,139,882 | 0.0101 | -0.232 | mu | R | 10q26.11 | chr10:118,307,938 | 0.0205 | -3.331 | tf | A | 3q13.33 | chr3:121,928,620 | 0.0293 | 1.2557 | tau | A | 12q24.13 | chr12:113,095,541 | 0.0348 | 2.609 | entropy | R | 12p11.23 | chr12:27,082,066 | 0.0403 | -0.104 | tf | A | 11p15.4 | chr11:8,683,933 | 0.0476 | 1.63 |
| tau | R | 3q21.2 | chr3:124,732,437 | 0.0015 | 7.275 | entropy | R | 5q23.1 | chr5:119,541,871 | 0.0101 | -0.292 | mu | R | 12q13.11 | chr12:47,748,560 | 0.0205 | 4.4422 | tf | A | 4q32.3 | chr4:165,484,648 | 0.0293 | 2.3674 | tau | A | 16p13.3 | chr16:3,056,002 | 0.0348 | 4.0806 | tau | R | 17q23.3 | chr17:63,929,637 | 0.0403 | 6.4078 | tf | A | 4q32.3 | chr4:165,340,437 | 0.0476 | 2.6774 |
| mu | R | 16p11.2 | chr16:28,592,334 | 0.0016 | 4.305 | entropy | R | 5q13.3 | chr5:76,632,129 | 0.0101 | -0.179 | tf | R | 16p13.13 | chr16:11,678,357 | 0.0205 | 5.3166 | entropy | D | 4p15.2 | chr4:25,124,081 | 0.0293 | -0.1394 | tf | R | 19p12 | chr19:21,635,190 | 0.0348 | 5.5555 | mu | A | 12q23.3 | chr12:104,326,306 | 0.0404 | 3.2576 | tau | A | 6q24.3 | chr6:146,726,205 | 0.0477 | 2.1189 |
| mu | R | 17q12 | chr17:39,451,343 | 0.0016 | 7.378 | entropy | R | 6p21.32 | chr6:33,432,001 | 0.0101 | -0.234 | mu | R | 1p22.1 | chr1:91,535,577 | 0.0205 | 3.0287 | tf | A | 5q35.2 | chr5:173,607,986 | 0.0293 | 2.2223 | tau | A | 19q13.43 | chr19:56,180,543 | 0.0348 | 2.7842 | tf | A | 11p15.4 | chr11:5,754,558 | 0.0407 | 1.7672 | tf | R | 20q13.33 | chr20:62,202,099 | 0.0478 | -1.846 |
| mu | R | 1p36.22 | chr1:9,267,354 | 0.0016 | 6.724 | mu | R | 18q21.2 | chr18:55,234,376 | 0.0102 | 5.744 | tf | R | 2q12.1 | chr2:105,089,049 | 0.0205 | 3.5668 | tf | A | 6q13 | chr6:72,235,789 | 0.0293 | 1.301 | tau | A | 2q14.3 | chr2:127,588,385 | 0.0348 | -2.011 | s | R | 16p13.12 | chr16:12,527,319 | 0.0407 | 0.491 | tf | D | 6p21.2 | chr6:38,989,995 | 0.0478 | -2.008 |
| tf | R | 4p12 | chr4:48,071,484 | 0.0017 | 7.446 | mu | R | 7q21.3 | chr7:94,413,927 | 0.0102 | 4.298 | mu | R | 2q24.3 | chr2:166,436,372 | 0.0205 | 4.2918 | entropy | D | 7p12.1 | chr7:53,003,071 | 0.0293 | -0.1558 | tau | A | 6q14.1 | chr6:78,978,649 | 0.0348 | 5.6357 | s | R | 19q13.12 | chr19:36,991,864 | 0.0407 | 0.6655 | mu | R | 15q26.3 | chr15:100,054,066 | 0.0479 | -4.366 |
| tf | R | 5p15.2 | chr5:10,239,149 | 0.0017 | 10.85 | mu | R | 16q24.3 | chr16:88,717,706 | 0.0104 | 2.897 | mu | R | 7p14.1 | chr7:37,907,356 | 0.0205 | 5.1221 | tf | A | 9q34.2 | chr9:133,317,737 | 0.0293 | 2.2169 | tau | R | 11p15.4 | chr11:4,368,357 | 0.0351 | -3.383 | s | R | 1p36.11 | chr1:26,324,059 | 0.0407 | 0.3003 | mu | R | 1p31.1 | chr1:78,629,896 | 0.0479 | 4.3646 |
| mu | R | 13q13.3 | chr13:39,013,029 | 0.0018 | 7.972 | mu | R | 20q13.32 | chr20:58,995,513 | 0.0104 | 4.329 | mu | R | 8p23.3 | chr8:468,993 | 0.0206 | 4.3571 | tf | A | 9p13.3 | chr9:34,981,984 | 0.0293 | -1.2807 | tau | R | 12q24.23 | chr12:119,630,059 | 0.0351 | 11.154 | s | R | 1p34.1 | chr1:45,620,657 | 0.0407 | -0.617 | s | A | 13q14.11 | chr13:42,302,154 | 0.0482 | 0.3582 |
| mu | R | 5q31.1 | chr5:133,993,134 | 0.0018 | 7.361 | entropy | R | 11q11 | chr11:55,796,424 | 0.0105 | -0.169 | tau | R | 1q25.1 | chr1:175,014,497 | 0.0207 | 5.3966 | mu | R | 3q27.3 | chr3:187,371,024 | 0.0295 | 2.7593 | tau | R | 14q23.3 | chr14:64,533,252 | 0.0351 | 6.4689 | s | R | 2q37.3 | chr2:239,297,974 | 0.0407 | 0.7387 | s | A | 16p12.3 | chr16:17,096,183 | 0.0482 | 0.1854 |
| mu | R | 6p22.1 | chr6:30,057,402 | 0.0018 | 7.587 | mu | R | 15q21.2 | chr15:52,192,644 | 0.0105 | 4.555 | entropy | R | 7p11.2 | chr7:56,639,966 | 0.0207 | -0.271 | tf | R | 9q33.2 | chr9:121,284,413 | 0.0295 | 4.8474 | tau | R | 16q22.2 | chr16:71,758,793 | 0.0351 | 4.6821 | s | R | 4q26 | chr4:119,396,394 | 0.0407 | 0.5626 | s | A | 16q23.3 | chr16:81,869,385 | 0.0482 | 0.2173 |
| tau | R | 11q13.1 | chr11:64,284,975 | 0.002 | 5.122 | mu | R | 8p22 | chr8:17,298,884 | 0.0105 | 6.316 | tf | R | 14q32.31 | chr14:100,967,132 | 0.0208 | 5.4018 | tau | A | 10q11.21 | chr10:42,830,216 | 0.0297 | 3.8859 | tau | R | 2q13 | chr2:111,165,807 | 0.0351 | 11.154 | s | R | 7q22.1 | chr7:100,574,461 | 0.0407 | 0.8639 | s | A | 17q11.2 | chr17:31,532,446 | 0.0482 | 0.3513 |
| tau | R | 1p36.21 | chr1:15,324,566 | 0.002 | 5.893 | tf | R | 19q13.2 | chr19:40,877,248 | 0.0106 | 3.793 | tf | R | 10q11.21 | chr10:44,373,408 | 0.0209 | 6.1464 | tf | A | 10q11.21 | chr10:45,443,298 | 0.0297 | 1.5648 | tau | R | 5q23.1 | chr5:119,393,258 | 0.0351 | 5.1212 | s | R | 7q36.1 | chr7:148,417,773 | 0.0407 | -0.338 | s | A | 2q24.2 | chr2:159,743,206 | 0.0482 | 0.4198 |
| tau | R | 1q23.1 | chr1:158,524,626 | 0.0021 | 10.44 | tau | R | 6p21.1 | chr6:41,795,070 | 0.0106 | 6.316 | tf | R | 6p22.1 | chr6:29,657,224 | 0.021 | 6.4236 | tf | A | 11q24.3 | chr11:130,877,49 | 0.0297 | 1.1127 | tau | R | 6q23.2 | chr6:134,259,980 | 0.0351 | 4.8087 | s | R | 9p13.3 | chr9:34,311,022 | 0.0407 | 0.8782 | s | A | 2p12 | chr2:79,026,975 | 0.0482 | -0.192 |
| entropy | R | 9q34.3 | chr9:136,799,144 | 0.0021 | -0.266 | mu | R | 12p13.32 | chr12:4,635,141 | 0.0107 | 5.234 | tf | R | 7q36.1 | chr7:150,572,454 | 0.021 | 5.2732 | tf | A | 13q12.12 | chr13:24,691,133 | 0.0297 | 2.5388 | tau | R | 6q13 | chr6:70,559,756 | 0.0351 | 11.154 | entropy | R | 11p11.2 | chr11:43,886,554 | 0.0409 | -0.129 | s | A | 4q21.22 | chr4:82,490,044 | 0.0482 | 0.2237 |
| tf | R | 15q26.1 | chr15:89,314,467 | 0.0022 | 3.091 | mu | R | 19q13.41 | chr19:51,592,307 | 0.0107 | 6.415 | tf | R | 7q36.1 | chr7:150,627,069 | 0.021 | 5.2732 | tf | A | 14q24.1 | chr14:67,677,706 | 0.0297 | 2.6046 | tau | R | 7q36.1 | chr7:150,742,955 | 0.0351 | 4.2437 | entropy | R | 7p22.3 | chr7:850,221 | 0.0409 | -0.258 | entropy | A | 16q24.2 | chr16:87,701,921 | 0.0483 | -0.082 |
| entropy | R | 6q22.1 | chr6:117,561,046 | 0.0022 | -0.264 | mu | R | 2q37.3 | chr2:241,768,770 | 0.0107 | 4.927 | mu | R | 22q11.21 | chr22:20,994,625 | 0.0212 | 5.5898 | tf | A | 15q26.1 | chr15:89,263,479 | 0.0297 | 1.1779 | entropy | D | 10p14 | chr10:7,381,998 | 0.0353 | -0.196 | mu | R | 11q14.2 | chr11:87,178,579 | 0.0411 | -4.863 | entropy | A | 19p12 | chr19:23,362,448 | 0.0483 | 0.0588 |
| mu | R | 2p16.1 | chr2:55,881,747 | 0.0023 | 7.334 | mu | R | 6p21.33 | chr6:31,538,914 | 0.0108 | 5.989 | mu | R | 3p21.31 | chr3:46,436,335 | 0.0213 | 4.8226 | tf | A | 16p12.1 | chr16:24,827,667 | 0.0297 | 1.4753 | entropy | D | 6p21.32 | chr6:32,198,607 | 0.0353 | -0.171 | tf | R | 12q23.1 | chr12:96,592,451 | 0.0411 | 3.5195 | tf | R | 1p36.12 | chr1:21,474,853 | 0.0483 | 4.237 |
| mu | R | 4q21.1 | chr4:76,169,853 | 0.0023 | 7.19 | s | D | 6p21.1 | chr6:43,222,397 | 0.0108 | 0.512 | tau | R | 16p12.1 | chr16:27,221,530 | 0.0214 | 3.2319 | tf | A | 17q21.33 | chr17:50,112,554 | 0.0297 | 1.8821 | tau | R | 22q11.22 | chr22:22,162,743 | 0.0354 | 6.8105 | entropy | R | 17p11.2 | chr17:18,141,579 | 0.0411 | -0.25 | entropy | A | 6p21.33 | chr6:31,139,871 | 0.0483 | -0.085 |
| tau | R | 1q24.2 | chr1:167,819,958 | 0.0024 | 13.53 | mu | R | 16p13.13 | chr16:11,461,160 | 0.0113 | 4.024 | tau | R | 19q13.11 | chr19:34,194,712 | 0.0214 | 7.3069 | tau | A | 19p13.3 | chr19:3,633,117 | 0.0297 | 4.0564 | tau | R | 2p23.3 | chr2:26,527,777 | 0.0354 | 6.0805 | tau | A | 17q21.2 | chr17:41,377,775 | 0.0411 | 3.783 | entropy | A | 8p11.21 | chr8:42,169,450 | 0.0483 | -0.117 |
| tau | R | 19q13.42 | chr19:54,173,409 | 0.0026 | 9.853 | mu | R | 5q35.2 | chr5:176,657,893 | 0.0113 | 5.219 | tau | R | 6q14.3 | chr6:85,287,424 | 0.0214 | 7.2665 | tf | A | 19q13.31 | chr19:43,744,726 | 0.0297 | -1.4371 | mu | R | 13q32.1 | chr13:95,887,950 | 0.0356 | 2.4304 | mu | R | 18q11.2 | chr18:22,997,706 | 0.0411 | 4.6153 | tau | R | 19p13.2 | chr19:7,931,021 | 0.0484 | 4.741 |
| tau | R | 11q24.2 | chr11:124,185,670 | 0.0027 | 4.148 | mu | R | 17q21.2 | chr17:42,307,544 | 0.0115 | 5.315 | tf | R | 11q23.3 | chr11:120,982,252 | 0.0218 | 4.2753 | tf | A | 20p12.1 | chr20:15,893,582 | 0.0297 | 2.2275 | mu | R | 15q22.2 | chr15:62,247,582 | 0.0356 | 2.0836 | tau | A | 18p11.32 | chr18:335,099 | 0.0411 | 3.7628 | tf | A | 10q24.2 | chr10:98,423,813 | 0.0487 | 1.7208 |
| tau | R | 6q24.3 | chr6:146,745,957 | 0.0028 | 6.994 | mu | R | 8q22.1 | chr8:96,251,190 | 0.0116 | 2.066 | tf | R | 14q11.2 | chr14:22,163,321 | 0.0218 | 3.7345 | tf | R | 20p13 | chr20:1,708,198 | 0.0297 | 2.0275 | mu | R | 5q12.3 | chr5:64,695,685 | 0.0356 | 4.0727 | mu | R | 19q13.33 | chr19:50,819,976 | 0.0411 | 3.1794 | tf | A | 12q24.31 | chr12:124,952,304 | 0.0487 | -1.127 |
| tau | D | 10q11.21 | chr10:42,747,509 | 0.0029 | 5.94 | entropy | R | 12p12.1 | chr12:24,814,406 | 0.0117 | -0.282 | tf | R | 15q26.1 | chr15:89,263,479 | 0.0218 | 2.3532 | tf | A | 2q11.2 | chr2:100,279,648 | 0.0297 | 1.421 | mu | A | 12q23.2 | chr12:102,917,009 | 0.0359 | 1.3814 | tf | R | 2q33.3 | chr2:206,959,824 | 0.0411 | 4.3582 | tf | D | 15q25.3 | chr15:85,734,943 | 0.0487 | 2.2734 |
| mu | R | 11q13.2 | chr11:68,025,980 | 0.0029 | 6.474 | entropy | R | 20p13 | chr20:1,648,960 | 0.0117 | -0.123 | tf | R | 16q24.1 | chr16:84,733,364 | 0.0218 | 6.211 | tau | A | 2q14.3 | chr2:127,635,686 | 0.0297 | -2.5639 | mu | A | 21q22.3 | chr21:44,558,709 | 0.0359 | 1.4484 | entropy | R | 6p21.31 | chr6:36,489,577 | 0.0411 | -0.143 | tf | A | 15q26.1 | chr15:89,319,135 | 0.0487 | 1.1295 |
| mu | R | 21q22.3 | chr21:44,558,704 | 0.0029 | 3.399 | tau | A | 3q12.2 | chr3:100,451,709 | 0.0117 | 2.943 | tf | R | 1q41 | chr1:219,988,517 | 0.0218 | 5.7936 | tau | A | 2q14.3 | chr2:127,637,380 | 0.0297 | 2.8569 | mu | A | 6p21.32 | chr6:32,197,092 | 0.0359 | 1.2273 | tau | R | 12q23.3 | chr12:106,311,196 | 0.0412 | 5.0641 | tf | A | 1q44 | chr1:248,294,677 | 0.0487 | 1.376 |
| tau | R | 5q22.2 | chr5:113,433,177 | 0.003 | 9.866 | mu | R | 11p11.2 | chr11:48,217,328 | 0.0118 | 5.225 | tf | R | 22q13.2 | chr22:42,794,643 | 0.0218 | -2.062 | tau | A | 2p15 | chr2:62,943,786 | 0.0297 | 3.8387 | mu | A | 7p12.3 | chr7:47,303,485 | 0.0359 | 2.1526 | tf | D | 14q24.1 | chr14:67,677,706 | 0.0412 | 2.6046 | tf | A | 20p12.2 | chr20:10,407,858 | 0.0487 | 2.0316 |
| tau | R | 8q24.11 | chr8:116,770,146 | 0.003 | 11.08 | entropy | R | 6q14.2 | chr6:83,665,658 | 0.0119 | -0.236 | tf | R | 5q15 | chr5:95,900,711 | 0.0218 | 3.9824 | tf | A | 4p14 | chr4:36,316,210 | 0.0297 | 3.0365 | mu | R | 13q12.11 | chr13:20,139,512 | 0.036 | 4.5324 | tf | D | 1q24.1 | chr1:166,852,946 | 0.0412 | -2.097 | tf | A | 6q23.3 | chr6:137,219,233 | 0.0487 | 1.1877 |
| tau | D | 16p13.3 | chr16:2,317,335 | 0.0031 | 5.087 | entropy | R | 8p21.1 | chr8:28,568,746 | 0.0119 | -0.124 | tf | R | 9p23 | chr9:13,176,313 | 0.0219 | 4.3782 | tau | A | 6p21.33 | chr6:31,364,274 | 0.0297 | 3.1567 | mu | R | 10p15.1 | chr10:6,464,519 | 0.0361 | 5.1187 | entropy | R | 2q14.3 | chr2:128,178,426 | 0.0412 | -0.097 | tf | A | 6p21.32 | chr6:32,829,532 | 0.0487 | 2.8637 |
| tau | R | 5q15 | chr5:94,714,683 | 0.0031 | 11.64 | tf | R | 14q24.3 | chr14:75,124,143 | 0.0121 | 2.417 | tf | R | 13q14.11 | chr13:42,189,018 | 0.022 | 4.1941 | tf | A | 6p21.2 | chr6:39,012,793 | 0.0297 | 2.4392 | mu | R | 11q24.1 | chr11:122,811,403 | 0.0361 | 3.8599 | tf | D | 5q14.3 | chr5:83,538,811 | 0.0412 | -2.105 | tf | D | 7q36.1 | chr7:150,802,179 | 0.0487 | -2.026 |
| tau | R | 10q22.3 | chr10:80,253,363 | 0.0032 | 8.651 | mu | R | 17p11.2 | chr17:16,939,677 | 0.0121 | 5.899 | mu | R | 1p13.3 | chr1:109,274,570 | 0.022 | 4.089 | tf | A | 7q36.1 | chr7:150,512,210 | 0.0297 | 1.307 | mu | R | 11p14.2 | chr11:26,670,934 | 0.0361 | 2.9229 | tau | R | 7q36.1 | chr7:149,126,274 | 0.0412 | 7.3802 | tf | D | 8q13.1 | chr8:65,605,251 | 0.0487 | 1.6012 |
| mu | R | 1p36.32 | chr1:3,412,095 | 0.0032 | 7.069 | tau | R | 2q14.3 | chr2:127,636,225 | 0.0121 | 7.523 | tau | R | 2p25.1 | chr2:7,041,347 | 0.0222 | 10.876 | tf | A | 7q36.3 | chr7:159,109,876 | 0.0297 | 2.1679 | mu | R | 12q24.33 | chr12:132,730,341 | 0.0361 | 1.8438 | tau | R | 7q11.23 | chr7:76,502,378 | 0.0412 | 3.993 | tf | A | 9q34.3 | chr9:137,612,856 | 0.0487 | 1.7346 |
| mu | R | 10q26.11 | chr10:119,669,744 | 0.0036 | -6.712 | tau | R | 3q27.1 | chr3:184,258,315 | 0.0121 | 9.89 | tf | R | 15q26.1 | chr15:90,360,270 | 0.0223 | 4.7332 | tf | A | 9q34.3 | chr9:137,588,887 | 0.0297 | 1.7207 | mu | R | 12p12.3 | chr12:18,648,442 | 0.0361 | -5.434 | mu | A | 11p15.5 | chr11:1,630,365 | 0.0417 | 2.3361 | tf | D | 2q33.1 | chr2:197,008,839 | 0.0488 | 2.0732 |
| tau | D | 12q23.3 | chr12:107,635,409 | 0.0036 | 4.366 | tf | R | 3p14.3 | chr3:58,410,164 | 0.0121 | 7.043 | tf | R | 2q24.2 | chr2:159,433,400 | 0.0223 | 5.9323 | tf | A | 9p13.3 | chr9:34,922,859 | 0.0297 | 1.2308 | mu | R | 19q13.33 | chr19:48,680,921 | 0.0361 | -5.113 | entropy | R | 14q11.2 | chr14:22,163,782 | 0.0417 | -0.151 | entropy | R | 3p25.2 | chr3:12,737,910 | 0.0488 | -0.155 |
| entropy | R | 16q24.2 | chr16:87,701,921 | 0.0036 | -0.282 | tau | R | 5q21.1 | chr5:103,086,739 | 0.0121 | 8.666 | mu | R | 10q24.1 | chr10:96,626,676 | 0.0225 | -5.542 | mu | R | 1p35.3 | chr1:29,203,187 | 0.0298 | 4.7999 | mu | R | 2q14.2 | chr2:119,441,564 | 0.0361 | 4.757 | tau | R | 16q22.1 | chr16:68,364,959 | 0.0417 | 8.4599 | entropy | R | 4p14 | chr4:38,021,681 | 0.0488 | -0.234 |
| mu | R | 18q23 | chr18:78,993,768 | 0.0036 | 5.166 | mu | R | 5q23.2 | chr5:123,378,295 | 0.0121 | 3.693 | mu | R | 11q13.1 | chr11:65,455,481 | 0.0225 | -2.55 | mu | R | 3p26.1 | chr3:5,216,746 | 0.0298 | 4.9505 | mu | R | 2q37.1 | chr2:233,294,901 | 0.0361 | 4.7676 | tau | R | 17q21.31 | chr17:44,910,155 | 0.0417 | 8.6564 | tf | A | 4q21.3 | chr4:86,015,955 | 0.0488 | 1.2825 |
| entropy | R | 4p15.1 | chr4:34,926,388 | 0.0036 | -0.324 | mu | R | 5q13.2 | chr5:69,123,227 | 0.0121 | 2.287 | mu | R | 19q13.2 | chr19:39,459,124 | 0.0225 | -2.218 | mu | R | 5q35.2 | chr5:173,961,094 | 0.0298 | 2.7802 | mu | R | 5q15 | chr5:96,795,133 | 0.0361 | 3.7375 | mu | A | 19p13.2 | chr19:11,832,882 | 0.0417 | -3.176 | entropy | R | 5q23.1 | chr5:119,393,258 | 0.0488 | -0.137 |
| mu | R | 6p21.32 | chr6:32,189,862 | 0.0036 | 5.798 | tf | A | 15q26.1 | chr15:89,314,467 | 0.0123 | 1.519 | mu | R | 20q13.32 | chr20:58,367,026 | 0.0225 | 2.3494 | tf | R | 5q31.1 | chr5:135,834,969 | 0.03 | 3.563 | mu | R | 7q31.2 | chr7:116,252,241 | 0.0361 | 5.4663 | mu | A | 20q11.22 | chr20:35,731,421 | 0.0417 | 2.1015 | entropy | R | 8q13.1 | chr8:66,453,625 | 0.0488 | -0.16 |
| mu | R | 7p14.3 | chr7:29,513,239 | 0.0036 | 6.901 | tau | R | 5q14.1 | chr5:79,791,060 | 0.0123 | 7.288 | tf | D | 2q32.1 | chr2:186,668,641 | 0.023 | -1.83 | entropy | A | 14q13.1 | chr14:33,214,249 | 0.0301 | -0.0878 | tf | A | 9q31.3 | chr9:110,385,959 | 0.0361 | 2.6456 | mu | A | 21q22.3 | chr21:44,305,738 | 0.0417 | 2.9751 | entropy | R | 9q34.3 | chr9:136,358,616 | 0.0488 | -0.295 |
| mu | R | 16p13.3 | chr16:792,706 | 0.0037 | 5.399 | tau | R | 6p21.2 | chr6:37,284,434 | 0.0123 | 4.612 | tf | R | 10q24.2 | chr10:99,685,834 | 0.0231 | 3.9566 | mu | R | 11p15.4 | chr11:4,164,233 | 0.0302 | 1.7891 | s | R | 16p13.3 | chr16:3,561,609 | 0.0362 | 0.321 | tau | R | 2q37.3 | chr2:240,467,232 | 0.0417 | 5.1272 | entropy | R | 9q34.3 | chr9:136,378,836 | 0.0488 | -0.295 |
| tf | R | 15q21.3 | chr15:58,640,733 | 0.0038 | 6.794 | mu | R | 2p11.2 | chr2:89,010,265 | 0.0125 | 4.618 | tf | R | 10p13 | chr10:12,832,283 | 0.0231 | 5.9925 | mu | R | 15q21.3 | chr15:58,640,733 | 0.0303 | 4.6849 | s | R | 9q33.3 | chr9:125,232,852 | 0.0362 | 0.8959 | tau | R | 2p16.1 | chr2:55,222,182 | 0.0417 | 7.3591 | tf | R | 10q26.13 | chr10:124,042,293 | 0.049 | -2.373 |
| tf | R | 1q23.3 | chr1:162,852,750 | 0.0038 | 4.005 | mu | R | 5q31.3 | chr5:141,661,868 | 0.0125 | 2.533 | tf | R | 19p13.2 | chr19:8,125,866 | 0.0231 | 4.7768 | tf | A | 15q25.3 | chr15:85,734,943 | 0.0303 | 2.2734 | tf | D | 10p12.1 | chr10:27,208,262 | 0.0364 | -2.791 | tau | R | 3q13.2 | chr3:112,184,615 | 0.0417 | 3.1822 | tf | A | 11p15.3 | chr11:12,940,713 | 0.049 | 1.2071 |
| tf | R | 1q41 | chr1:223,771,750 | 0.0038 | 7.466 | mu | R | 8q24.3 | chr8:142,736,388 | 0.0126 | 4.404 | tf | R | 3q13.33 | chr3:121,922,681 | 0.0231 | 2.2156 | mu | R | 17q21.2 | chr17:42,313,598 | 0.0303 | 4.1257 | tf | D | 11q23.1 | chr11:110,712,111 | 0.0364 | 2.622 | mu | A | 3q26.1 | chr3:167,789,149 | 0.0417 | 2.121 | tf | A | 11q13.3 | chr11:68,980,016 | 0.049 | 2.7324 |
| tau | R | 5q23.1 | chr5:119,527,040 | 0.0038 | 9.384 | tau | D | 11p15.4 | chr11:6,151,770 | 0.0127 | 5.738 | tf | R | 3q27.2 | chr3:185,048,513 | 0.0231 | 4.0413 | mu | R | 20q11.23 | chr20:38,649,915 | 0.0303 | 4.3662 | tf | D | 12q24.31 | chr12:124,952,304 | 0.0364 | -2.024 | mu | A | 4q31.1 | chr4:138,561,804 | 0.0417 | 1.4385 | tf | A | 12q23.3 | chr12:104,326,306 | 0.049 | 2.8129 |
| tau | R | 7p22.3 | chr7:290,856 | 0.0038 | 7.905 | tau | D | 1p36.23 | chr1:7,781,270 | 0.0127 | 5.033 | tf | R | 5q33.2 | chr5:155,015,898 | 0.0231 | 4.8721 | tf | A | 21q22.3 | chr21:46,431,839 | 0.0303 | 1.4856 | tf | D | 16q24.1 | chr16:84,179,247 | 0.0364 | 3.3662 | mu | A | 4q32.3 | chr4:169,007,013 | 0.0417 | 1.9509 | tf | A | 16p13.3 | chr16:2,853,080 | 0.049 | 1.2353 |
| mu | R | 16q23.1 | chr16:74,716,498 | 0.0041 | 6.836 | tau | R | 13q12.13 | chr13:24,951,343 | 0.0128 | 6.283 | tf | R | 4p16.3 | chr4:3,485,510 | 0.0232 | 3.3101 | tf | A | 2q33.1 | chr2:197,008,839 | 0.0303 | 2.0732 | tf | D | 19p13.2 | chr19:11,222,682 | 0.0364 | 2.9659 | tau | R | 6p22.3 | chr6:18,148,016 | 0.0417 | 3.3684 | tf | A | 17q12 | chr17:35,650,519 | 0.049 | -1.95 |
| entropy | D | 19p12 | chr19:23,362,448 | 0.0041 | 0.119 | tau | R | 20q11.21 | chr20:32,997,626 | 0.0128 | 9.942 | tau | R | 5q33.1 | chr5:151,491,042 | 0.0232 | 7.3171 | mu | R | 2q37.3 | chr2:240,552,838 | 0.0303 | 2.7851 | tf | D | 19p13.2 | chr19:12,440,915 | 0.0364 | 3.0704 | mu | A | 7p12.3 | chr7:45,714,555 | 0.0417 | 1.8426 | tf | A | 19q13.33 | chr19:49,877,601 | 0.049 | -1.066 |
| entropy | D | 1q24.2 | chr1:169,703,202 | 0.0041 | -0.123 | tau | R | 6p21.1 | chr6:42,465,630 | 0.0128 | 9.104 | s | R | 2q36.2 | chr2:224,511,315 | 0.0234 | 0.696 | tf | A | 3q29 | chr3:195,774,278 | 0.0303 | -1.149 | entropy | R | 1q23.1 | chr1:159,015,862 | 0.0364 | -0.269 | tf | A | 10p11.23 | chr10:30,611,750 | 0.0419 | 1.9887 | tf | A | 19q13.33 | chr19:49,880,379 | 0.049 | -1.111 |
| mu | R | 1q32.1 | chr1:205,075,226 | 0.0041 | 5.82 | tau | R | 6q14.3 | chr6:85,489,515 | 0.0128 | 6.66 | s | R | 7q36.3 | chr7:158,926,393 | 0.0234 | 0.6853 | mu | R | 6q22.1 | chr6:117,602,436 | 0.0303 | -1.9905 | tf | D | 2p24.1 | chr2:20,239,735 | 0.0364 | 3.1503 | s | R | 14q11.2 | chr14:22,770,303 | 0.0419 | 0.3681 | tf | A | 1q21.3 | chr1:154,378,113 | 0.049 | 1.5185 |
| mu | R | 1q43 | chr1:241,633,141 | 0.0041 | 5.739 | tau | R | 11q24.2 | chr11:124,146,384 | 0.0131 | 5.014 | tf | R | 11p15.5 | chr11:2,308,765 | 0.0236 | 5.4085 | mu | A | 11p15.1 | chr11:20,637,133 | 0.0305 | -1.6623 | tf | D | 3p25.3 | chr3:10,002,578 | 0.0364 | -1.995 | tf | A | 15q25.2 | chr15:82,573,126 | 0.0419 | 2.4536 | tf | A | 1q23.3 | chr1:162,855,468 | 0.049 | 1.5723 |
| mu | R | 2q11.2 | chr2:101,281,950 | 0.0041 | 5.549 | entropy | R | 21q22.3 | chr21:43,791,051 | 0.0131 | -0.221 | tf | R | 17q21.33 | chr17:50,112,554 | 0.0236 | 5.9458 | mu | A | 11p11.2 | chr11:43,886,680 | 0.0305 | 2.2374 | tf | D | 3q29 | chr3:195,774,278 | 0.0364 | -1.735 | tf | A | 19p13.3 | chr19:4,210,359 | 0.0419 | 1.2495 | tf | A | 20p13 | chr20:275,932 | 0.049 | -1.214 |
| mu | R | 4p12 | chr4:48,071,484 | 0.0041 | 5.96 | s | R | 3q13.33 | chr3:119,517,170 | 0.0134 | 0.565 | mu | R | 2q21.3 | chr2:135,207,806 | 0.0238 | 4.4002 | mu | A | 12q24.33 | chr12:132,675,97 | 0.0305 | 1.4935 | tf | D | 9q34.2 | chr9:133,317,737 | 0.0364 | 2.4636 | tf | A | 22q13.2 | chr22:42,794,643 | 0.0419 | -1.168 | tau | R | 3q25.1 | chr3:149,474,645 | 0.049 | 6.5861 |
| mu | R | 4p16.3 | chr4:654,920 | 0.0041 | 6.269 | mu | R | 3p25.1 | chr3:16,271,813 | 0.0134 | 6.428 | entropy | R | 17p12 | chr17:11,929,970 | 0.0239 | -0.122 | mu | A | 1q24.2 | chr1:167,789,761 | 0.0305 | 2.6556 | entropy | R | 15q25.1 | chr15:78,998,007 | 0.0367 | -0.207 | tf | A | 2q32.1 | chr2:186,668,641 | 0.0419 | -1.033 | tf | A | 3q27.2 | chr3:185,191,137 | 0.049 | 2.3653 |
| mu | R | 4q22.1 | chr4:88,707,091 | 0.0041 | 6.66 | mu | R | 6p12.3 | chr6:47,899,741 | 0.0135 | 3.971 | entropy | R | 19q13.12 | chr19:35,812,019 | 0.0239 | -0.153 | tf | R | 12q12 | chr12:44,780,101 | 0.0306 | 4.2993 | mu | R | 17p13.3 | chr17:2,800,613 | 0.0367 | 5.4363 | tf | A | 3q13.33 | chr3:121,844,438 | 0.0419 | 1.1853 | tf | A | 3p14.1 | chr3:64,013,646 | 0.049 | -1.467 |
| mu | R | 6p22.1 | chr6:30,108,066 | 0.0041 | 6.186 | tf | R | 11p15.1 | chr11:17,510,555 | 0.0136 | 6.898 | entropy | R | 22q12.3 | chr22:32,475,396 | 0.024 | -0.212 | tf | A | 14q24.3 | chr14:73,714,341 | 0.0306 | 1.5236 | mu | R | 3q13.33 | chr3:120,324,989 | 0.0367 | 4.9018 | tf | A | 3q21.3 | chr3:126,130,395 | 0.0419 | -1.323 | tf | A | 3q12.1 | chr3:98,793,766 | 0.049 | 2.3216 |
| mu | R | 11q24.3 | chr11:129,873,117 | 0.0042 | 5.789 | tau | R | 10q26.2 | chr10:126,016,711 | 0.0137 | 4.57 | tau | R | 2q31.3 | chr2:181,482,345 | 0.024 | 9.2344 | tf | A | 17q11.2 | chr17:32,369,273 | 0.0306 | 1.3334 | mu | R | 3p14.3 | chr3:56,727,687 | 0.0367 | 4.6446 | tf | A | 3q23 | chr3:142,213,514 | 0.0419 | -1.389 | tf | A | 5q31.1 | chr5:135,834,969 | 0.049 | 1.3279 |
| mu | R | 12q23.2 | chr12:103,406,351 | 0.0042 | 5.044 | entropy | R | 6p12.3 | chr6:46,829,592 | 0.0137 | -0.209 | tau | D | 19q13.11 | chr19:32,879,255 | 0.0241 | -3.37 | tf | A | 19p13.2 | chr19:11,222,682 | 0.0306 | 2.482 | mu | R | 6p24.3 | chr6:7,310,026 | 0.0367 | -2.314 | tf | A | 6p24.2 | chr6:11,103,655 | 0.0419 | 1.4891 | tf | A | 5q14.3 | chr5:90,514,470 | 0.049 | 2.166 |
| mu | R | 14q32.12 | chr14:94,046,150 | 0.0042 | 7.386 | mu | R | 10p11.1 | chr10:38,645,999 | 0.0138 | 3.926 | entropy | R | 2q31.3 | chr2:179,945,453 | 0.0241 | -0.169 | tf | A | 21q22.3 | chr21:46,194,555 | 0.0306 | 1.599 | mu | R | 18q21.33 | chr18:62,442,927 | 0.0368 | 5.7988 | tf | A | 7q36.1 | chr7:150,802,179 | 0.0419 | -1.237 | tf | A | 6p21.33 | chr6:31,354,710 | 0.049 | 2.3682 |
| mu | R | 16p13.3 | chr16:1,457,842 | 0.0042 | 7.243 | mu | R | 17q21.31 | chr17:43,529,343 | 0.0138 | 5.199 | tau | D | 9q21.13 | chr9:76,194,730 | 0.0241 | 4.6114 | tf | A | 7p21.3 | chr7:7,474,604 | 0.0306 | -1.4176 | tau | R | 3p24.3 | chr3:17,012,007 | 0.0368 | 3.5024 | tau | A | 1q23.1 | chr1:157,045,172 | 0.0421 | -1.997 | tf | A | 6p21.33 | chr6:31,806,044 | 0.049 | 2.0534 |
| mu | R | 1q23.1 | chr1:156,914,792 | 0.0042 | 7.039 | mu | R | 17q25.3 | chr17:82,657,767 | 0.0139 | 5.253 | entropy | R | 16p12.1 | chr16:27,215,116 | 0.0245 | -0.096 | tf | R | 11q23.3 | chr11:118,120,68 | 0.031 | 1.3501 | entropy | R | 11q13.2 | chr11:68,042,874 | 0.0372 | -0.151 | tau | A | 3q21.2 | chr3:126,006,809 | 0.0421 | 3.3475 | tf | A | 11q24.2 | chr11:125,959,610 | 0.0491 | 1.9643 |

[illegible]

**Supplementary Table 2** Differentially expressed genes upon TBKBP1 overexpression in MM cell lines

| baseMean | log2FoldChange | lfcSE | stat | pvalue | padj | Ensembl |
|---|---|---|---|---|---|---|
| 65.7744928 | -1.093648505 | 0.28179912 | -3.8809508 | 0.00010405 | 0.00918683 | ENSG00000228526 |
| 6911.90077 | 1.070800132 | 0.25826989 | 4.14605104 | 3.382584500 | 0.00433706 | ENSG00000117450 |
| 38.9311763 | 1.308569846 | 0.39127287 | 3.34439193 | 0.00082463 | 0.03217769 | ENSG00000172260 |
| 98.2339432 | 1.793115889 | 0.45442876 | 3.94586798 | 7.951141655 | 0.00814164 | ENSG00000137965 |
| 56.5072248 | 1.55257252 | 0.37566272 | 4.13288948 | 3.582310006 | 0.00455355 | ENSG00000224093 |
| 34.9031339 | -1.076105479 | 0.31170895 | -3.4522765 | 0.00055588 | 0.02553401 | ENSG00000232811 |
| 238.200718 | -1.050688662 | 0.27756641 | -3.7853595 | 0.00015349 | 0.01172621 | ENSG00000184678 |
| 244.922826 | 1.114244293 | 0.26704966 | 4.17242352 | 3.013767707 | 0.00396768 | ENSG00000027869 |
| 1887.02336 | -1.498832673 | 0.29011885 | -5.1662713 | 2.388100983 | 0.0001531 | ENSG00000132185 |
| 40.3323082 | 1.285248794 | 0.3954426 | 3.25015262 | 0.00115343 | 0.03856539 | ENSG00000162745 |
| 41.6865086 | -1.652935215 | 0.44958436 | -3.6765852 | 0.00023638 | 0.01483139 | ENSG00000143341 |
| 40.3292611 | -1.521540279 | 0.42571271 | -3.5741011 | 0.00035143 | 0.01877493 | ENSG00000174307 |
| 25.1936182 | -1.584878717 | 0.39257597 | -4.0371262 | 5.410997575 | 0.00609047 | ENSG00000198892 |
| 55.5995963 | 1.587132117 | 0.41252666 | 3.84734432 | 0.00011941 | 0.01000357 | ENSG00000133048 |
| 392.315544 | 1.364410298 | 0.2076947 | 6.56930717 | 5.054990837 | 1.064797712 | ENSG00000136634 |
| 450.313738 | -1.123945834 | 0.21528545 | -5.2207235 | 1.782254580 | 0.00011945 | ENSG00000116991 |
| 27.2362698 | 1.414256628 | 0.42204329 | 3.35097523 | 0.00080528 | 0.03178268 | ENSG00000226398 |
| 30.5503526 | -1.458620401 | 0.37963302 | -3.8421853 | 0.00012194 | 0.01015853 | ENSG00000230002 |
| 436.529 | 1.330945791 | 0.26941277 | 4.9401734 | 7.805312723 | 0.00030287 | ENSG00000115085 |
| 68.8260169 | 1.183270394 | 0.36325372 | 3.2574213 | 0.00112429 | 0.03793529 | ENSG00000115526 |
| 209.374805 | -1.043603983 | 0.23659643 | -4.4109033 | 1.029402937 | 0.00185733 | ENSG00000182263 |
| 29.4123721 | -1.752643579 | 0.42793863 | -4.0955489 | 4.211693012 | 0.00508909 | ENSG00000136531 |
| 474.283567 | 1.350924616 | 0.26712188 | 5.05733412 | 4.251579673 | 0.0001959 | ENSG00000160791 |
| 148.960496 | -1.22538927 | 0.23999632 | -5.1058669 | 3.292814100 | 0.00018375 | ENSG00000080224 |
| 239.021543 | -1.057069929 | 0.28563477 | -3.7007747 | 0.00021494 | 0.01414876 | ENSG00000144821 |
| 283.184064 | 1.757423609 | 0.35025466 | 5.01755952 | 5.233199126 | 0.00023223 | ENSG00000163606 |
| 38.1109979 | -1.01642775 | 0.32345984 | -3.1423615 | 0.00167591 | 0.04489695 | ENSG00000286585 |
| 60.5043344 | -1.267617322 | 0.33389361 | -3.7964708 | 0.00014677 | 0.01151135 | ENSG00000085276 |
| 23.8210965 | -1.525451501 | 0.39377279 | -3.8739383 | 0.00010709 | 0.00928853 | ENSG00000171121 |
| 70.9787871 | -1.233141245 | 0.32980867 | -3.7389595 | 0.00018478 | 0.01285204 | ENSG00000242512 |
| 1463.29711 | -1.400128072 | 0.45076129 | -3.1061409 | 0.00189546 | 0.04822619 | ENSG00000078081 |
| 34.8356576 | -1.279274468 | 0.38493076 | -3.3233885 | 0.00088931 | 0.03335186 | ENSG00000188001 |
| 51.1500785 | -1.017093825 | 0.32189318 | -3.1597247 | 0.00157918 | 0.04373567 | ENSG00000305512 |
| 40.6717931 | -2.020404871 | 0.54643886 | -3.6974033 | 0.00021782 | 0.01427422 | ENSG00000251095 |
| 2251.0104 | -1.246245641 | 0.31638191 | -3.9390546 | 8.180330032 | 0.00826157 | ENSG00000170006 |
| 87.1581936 | 1.21224741 | 0.30340721 | 3.995447 | 6.457230219 | 0.00705273 | ENSG00000038295 |
| 139.948029 | 1.611588465 | 0.36678907 | 4.39377449 | 1.113993757 | 0.00192537 | ENSG00000137628 |
| 99.6238359 | 2.291591091 | 0.71699815 | 3.19609068 | 0.00139303 | 0.04141185 | ENSG00000181381 |
| 24.1186996 | -1.613974989 | 0.47424704 | -3.4032368 | 0.00066593 | 0.02854381 | ENSG00000225940 |
| 188.806219 | -1.05336252 | 0.28051216 | -3.7551404 | 0.00017324 | 0.012522 | ENSG00000197822 |
| 227.109291 | 1.597376253 | 0.36079462 | 4.42738376 | 9.538294421 | 0.00178807 | ENSG00000145685 |
| 109.621703 | 1.018704377 | 0.23391532 | 4.35501354 | 1.330586920 | 0.00217994 | ENSG00000233828 |
| 31.1882356 | -1.831081691 | 0.49271714 | -3.7162939 | 0.00020217 | 0.01354974 | ENSG00000145777 |
| 292.164942 | 1.538687361 | 0.23305718 | 6.60218813 | 4.051329147 | 9.956141378 | ENSG00000113070 |
| 278.578514 | 1.856169141 | 0.31776459 | 5.84133409 | 5.178443552 | 4.772259386 | ENSG00000135077 |
| 116.200095 | 1.061932882 | 0.33245095 | 3.19425434 | 0.00140193 | 0.04143531 | ENSG00000272142 |
| 64.9225654 | 2.043471824 | 0.59389881 | 3.4407744 | 0.00058005 | 0.02615555 | ENSG00000112799 |
| 99.364686 | -1.33192766 | 0.38989534 | -3.4161159 | 0.00063521 | 0.02760493 | ENSG00000124635 |
| 49.6897292 | -1.118904238 | 0.31400955 | -3.5632809 | 0.00036625 | 0.0192869 | ENSG00000206344 |
| 106.968056 | 1.218404729 | 0.30388094 | 4.00948058 | 6.085245829 | 0.00672242 | ENSG00000196735 |
| 67.5764117 | 4.078139316 | 0.42323497 | 9.63563888 | 5.653918603 | 4.168351490 | ENSG00000237541 |
| 251.776369 | 1.205261993 | 0.2707474 | 4.45161059 | 8.522862199 | 0.00165355 | ENSG00000156535 |
| 26.1029534 | -1.33524531 | 0.36051642 | -3.7037018 | 0.00021248 | 0.01411243 | ENSG00000132429 |
| 56.6630512 | -1.29646165 | 0.41929093 | -3.0920336 | 0.0019879 | 0.04974859 | ENSG00000105928 |
| 745.586392 | 1.386452763 | 0.28741981 | 4.82378994 | 1.408557658 | 0.00041538 | ENSG00000146674 |
| 101.702794 | -1.211496372 | 0.32154294 | -3.7677592 | 0.00016472 | 0.01232887 | ENSG00000165215 |
| 210.728787 | -1.004608683 | 0.23412682 | -4.290874 | 1.779712819 | 0.0025982 | ENSG00000105976 |
| 201.354029 | -1.001960137 | 0.20566516 | -4.8718029 | 1.105844641 | 0.00037861 | ENSG00000077063 |
| 57.3921408 | -1.766785648 | 0.54395266 | -3.2480504 | 0.00116199 | 0.03864055 | ENSG00000234722 |
| 46.4269276 | -2.100270062 | 0.58419093 | -3.5951775 | 0.00032417 | 0.01783543 | ENSG00000308970 |
| 3768.99225 | 4.679684855 | 0.89024319 | 5.2566365 | 1.467138912 | 0.00010816 | ENSG00000164690 |
| 69.8414379 | -1.085959407 | 0.32646672 | -3.3264016 | 0.00087975 | 0.03330034 | ENSG00000120896 |
| 76.3097688 | 1.045831919 | 0.31854581 | 3.28314444 | 0.00102656 | 0.0358688 | ENSG00000187955 |
| 319.655025 | 1.020366853 | 0.26248132 | 3.88738853 | 0.00010133 | 0.00905508 | ENSG00000286122 |
| 95.7169765 | 1.170981434 | 0.26443722 | 4.42820205 | 9.502186664 | 0.00178807 | ENSG00000184156 |
| 31.471396 | 2.26670387 | 0.70564158 | 3.21225951 | 0.00131695 | 0.0400381 | ENSG00000254338 |
| 84.7019133 | 3.542183178 | 0.38705514 | 9.15162411 | 5.608393016 | 2.067393875 | ENSG00000182759 |
| 29.4890542 | -1.485450433 | 0.3959144 | -3.7519484 | 0.00017547 | 0.01255942 | ENSG00000147862 |
| 332.34529 | -1.100682599 | 0.22612564 | -4.8675709 | 1.129783092 | 0.00037861 | ENSG00000147883 |
| 25.3338169 | -1.151182933 | 0.36233082 | -3.1771599 | 0.00148725 | 0.04280817 | ENSG00000226237 |
| 405.067306 | 1.161125492 | 0.27190957 | 4.27026338 | 1.952422630 | 0.002795 | ENSG00000136929 |
| 496.840673 | -1.091957405 | 0.33063952 | -3.3025617 | 0.00095806 | 0.03462401 | ENSG00000120594 |
| 135.358756 | -1.205006904 | 0.3430919 | -3.5121987 | 0.00044442 | 0.02224498 | ENSG00000078114 |
| 231.020442 | 2.869430304 | 0.74463428 | 3.85347599 | 0.00011645 | 0.00986836 | ENSG00000169129 |
| 77.1964883 | -1.272484371 | 0.38092397 | -3.3405206 | 0.00083621 | 0.03236217 | ENSG00000184545 |
| 323.195531 | -1.26403129 | 0.28812913 | -4.3870305 | 1.149086796 | 0.00192537 | ENSG00000132256 |
| 149.884155 | -1.130348638 | 0.3022278 | -3.7400551 | 0.00018398 | 0.01285204 | ENSG00000184014 |
| 24.8256011 | -1.364059473 | 0.41588871 | -3.2798665 | 0.00103856 | 0.03620236 | ENSG00000175868 |
| 52.9347766 | 1.29162449 | 0.37671269 | 3.42867266 | 0.00060654 | 0.02710134 | ENSG00000184937 |
| 116.848311 | 1.479313201 | 0.37822361 | 3.91121327 | 9.183363337 | 0.00873604 | ENSG00000179241 |
| 53.5664586 | -1.880328994 | 0.57919516 | -3.2464515 | 0.00116853 | 0.03864055 | ENSG00000148948 |
| 22.8278247 | -1.690353565 | 0.53828375 | -3.1402649 | 0.00168795 | 0.04500695 | ENSG00000110492 |
| 128.701455 | 1.041816256 | 0.32616008 | 3.19418688 | 0.00140225 | 0.04143531 | ENSG00000173621 |
| 33.2476287 | 1.488263858 | 0.46929322 | 3.17128776 | 0.00151765 | 0.04287291 | ENSG00000171631 |
| 32.9842273 | 1.581409321 | 0.43392823 | 3.64440292 | 0.00026801 | 0.01606445 | ENSG00000110237 |
| 95.676411 | 1.595349401 | 0.32631692 | 4.88895702 | 1.013716208 | 0.00036457 | ENSG00000137474 |
| 151.733492 | -1.336678342 | 0.41926587 | -3.1881401 | 0.00143191 | 0.04176497 | ENSG00000110675 |
| 136.58221 | 1.656729209 | 0.38547809 | 4.29785572 | 1.724583572 | 0.00256858 | ENSG00000255274 |
| 40.7958738 | -1.779947733 | 0.41664849 | -4.2720609 | 1.936746242 | 0.002795 | ENSG00000166257 |
| 205.17521 | 1.675222433 | 0.36839332 | 4.54737457 | 5.431927260 | 0.00119266 | ENSG00000171860 |
| 27.0084914 | 1.215914685 | 0.35959283 | 3.38136516 | 0.00072127 | 0.03004257 | ENSG00000139055 |
| 576.405736 | -1.245409489 | 0.19105758 | -6.5185033 | 7.101238746 | 1.308847066 | ENSG00000018236 |
| 33.121754 | -1.177863913 | 0.32708261 | -3.6011206 | 0.00031685 | 0.01768758 | ENSG00000275481 |
| 267.522519 | 1.033262198 | 0.29019513 | 3.56057735 | 0.00037004 | 0.01933014 | ENSG00000181418 |
| 849.891587 | 1.102868752 | 0.18685257 | 5.90234723 | 3.583659109 | 3.774360968 | ENSG00000139351 |
| 33.5754199 | 1.163934198 | 0.35122248 | 3.31395135 | 0.00091988 | 0.03384004 | ENSG00000255650 |
| 19.5868107 | -1.550535229 | 0.45606559 | -3.3998075 | 0.00067433 | 0.02873711 | ENSG00000151164 |
| 54.2188173 | -1.656574139 | 0.36563659 | -4.5306574 | 5.880043102 | 0.00125654 | ENSG00000198598 |
| 243.4416 | -1.190482522 | 0.29683799 | -4.0105464 | 6.057840606 | 0.00672242 | ENSG00000073910 |
| 49.7268379 | 1.84904589 | 0.42975968 | 4.30251129 | 1.688730660 | 0.00254085 | ENSG00000304344 |
| 419.098226 | 1.393673786 | 0.32926849 | 4.23263642 | 2.309677232 | 0.00315335 | ENSG00000198133 |
| 43.1464956 | -1.109793145 | 0.28635896 | -3.8755314 | 0.00010639 | 0.00928257 | ENSG00000070778 |
| 89.4490792 | 1.51953168 | 0.34627073 | 4.38827646 | 1.142524845 | 0.00192537 | ENSG00000119714 |
| 66.6610553 | 1.880331748 | 0.60554312 | 3.10519877 | 0.00190151 | 0.04825782 | ENSG00000182809 |
| 91.1959123 | -1.014670908 | 0.20028477 | -5.0661411 | 4.059610019 | 0.0001959 | ENSG00000285077 |
| 140.2694 | -1.087997218 | 0.33739466 | -3.224702 | 0.00126104 | 0.03941388 | ENSG00000129028 |
| 291.848872 | 2.251279464 | 0.34679096 | 6.49174784 | 8.484618843 | 1.390063387 | ENSG00000008517 |
| 73.6778595 | 1.476308582 | 0.28937134 | 5.10177891 | 3.364756871 | 0.00018375 | ENSG00000103534 |
| 22.7416866 | -1.661803635 | 0.5175462 | -3.2109281 | 0.00132307 | 0.0401413 | ENSG00000182601 |
| 231.899211 | 1.049419646 | 0.22508418 | 4.66234304 | 3.126295104 | 0.00075569 | ENSG00000167703 |
| 38.6834625 | -1.479421487 | 0.42598196 | -3.4729674 | 0.00051474 | 0.02413097 | ENSG00000141756 |
| 15251.0171 | 11.95590733 | 1.85314523 | 6.45168396 | 1.106140691 | 1.631004449 | ENSG00000198933 |
| 878.953419 | -1.139749623 | 0.12147047 | -9.3829359 | 6.416077309 | 3.153501997 | ENSG00000171298 |
| 69.399527 | 3.640897277 | 0.51692647 | 7.04335624 | 1.876636111 | 5.534199892 | ENSG00000181409 |
| 167.234029 | 2.111705369 | 0.41750652 | 5.05789796 | 4.239031454 | 0.0001959 | ENSG00000141574 |
| 66.4968499 | 2.752871412 | 0.54315582 | 5.06829033 | 4.014048238 | 0.0001959 | ENSG00000299255 |
| 37.5414917 | -1.715824473 | 0.38763067 | -4.4264415 | 9.580036441 | 0.00178807 | ENSG00000293697 |
| 96.7764417 | -1.126957543 | 0.34236651 | -3.2916699 | 0.00099594 | 0.03538603 | ENSG00000141449 |
| 153.721434 | -1.186026339 | 0.30337166 | -3.909483 | 9.249386141 | 0.00874245 | ENSG00000187323 |
| 490.546887 | -1.874541134 | 0.19430329 | -9.6475009 | 5.036823869 | 4.168351490 | ENSG00000179855 |
| 22.1692585 | -2.243553685 | 0.64503094 | -3.478211 | 0.00050477 | 0.02393206 | ENSG00000171124 |
| 36.4898724 | -1.12570181 | 0.32810725 | -3.4308958 | 0.00060159 | 0.02696191 | ENSG00000267169 |
| 148.059637 | -1.407079778 | 0.34371735 | -4.0937118 | 4.245219598 | 0.00508909 | ENSG00000072071 |
| 40.6579236 | 1.43057871 | 0.45534156 | 3.14177055 | 0.0016793 | 0.04489695 | ENSG00000260725 |
| 159.31533 | -1.007088818 | 0.27980189 | -3.5992924 | 0.00031908 | 0.01768758 | ENSG00000167608 |
| 61.4868464 | -1.155931164 | 0.35973413 | -3.213293 | 0.00131222 | 0.03997673 | ENSG00000239998 |
| 69.2897258 | 2.855927532 | 0.74678905 | 3.82427613 | 0.00013116 | 0.01072437 | ENSG00000101306 |
| 19.5264011 | -2.44632912 | 0.63579832 | -3.8476495 | 0.00011926 | 0.01000357 | ENSG00000026559 |
| 80.1213509 | -1.008256862 | 0.31256644 | -3.2257362 | 0.00125649 | 0.03941388 | ENSG00000215386 |
| 25.3995989 | -1.857627052 | 0.50707549 | -3.6634132 | 0.00024888 | 0.01535433 | ENSG00000159261 |
| 2204.44162 | 2.188908394 | 0.65827534 | 3.32521706 | 0.0008835 | 0.03331756 | ENSG00000183036 |
| 45.8446391 | -1.508224304 | 0.48325985 | -3.1209386 | 0.00180276 | 0.04696403 | ENSG00000211674 |
| 21.626938 | -2.686065364 | 0.70818939 | -3.7928631 | 0.00014892 | 0.01153635 | ENSG00000130487 |
| 1289.60855 | 1.059731952 | 0.28751085 | 3.68588506 | 0.00022791 | 0.01466114 | ENSG00000173198 |
